# Building Predictive Understanding of Microbial Ecology by Bridging Microbial Growth Kinetics and Microbial Population Dynamics

**DOI:** 10.1101/2024.07.25.605167

**Authors:** Zhang Cheng, Weibo Xia, Sean McKelvey, Qiang He, Yuzhou Chen, Heyang Yuan

**Author notes:** **Corresponding author** Yuzhou Chen, Heyang Yuan.

## Abstract

Modeling microbial communities can provide predictive insights into microbial ecology, but current modeling approaches suffer from inherent limitations. In this study, a novel modeling approach was proposed to address those limitations based on the intrinsic connection between the growth kinetics of guilds and the dynamics of individual microbial populations. To implement the modeling approach, 466 samples from four full-scale activated sludge systems were retrieved from the literature. The raw samples were processed using a data transformation method that not only increased the dataset size by three times but also enabled quantification of population dynamics. Most of the 42 family-level core populations showed overall dynamics close to zero within the sampling period, explaining their resilience to environmental perturbation. Bayesian networks built with environmental factors, perturbation, historical abundance, population dynamics, and mechanistically derived microbial kinetic parameters classified the core populations into heterotrophic and autotrophic guilds. Topological data analysis was applied to identify keystone populations and their time-dependent interactions with other populations. The data-driven inferences were validated directly using the Microbial Database for Activated Sludge (MiDAS) and indirectly by predicting population abundance and community structure using artificial neural networks. The Bray-Curtis similarity between predicted and observed communities was significantly higher with microbial kinetic parameters than without parameters (0.70 vs. 0.66), demonstrating the accuracy of the modeling approach. Implemented based on engineered systems, this modeling approach can be generalized to natural systems to gain predictive understandings of microbial ecology.

## 1. Introduction

Microorganisms play critical roles in various ecosystems by interacting with each other and forming microbial communities (Falkowski et al. 2008, Rittmann and McCarty 2012, Yatsunenko et al. 2012). How individual microorganisms function and interact within communities has always been a core question in microbial ecology (Coyte et al. 2015, Fuhrman 2009, Fuhrman et al. 2015, Stams and Plugge 2009, Torsvik and Øvreås 2002).

Our understanding of microbial ecology has been significantly advanced by the development of experimental and computational approaches (Ju and Zhang 2015b, Vanwonterghem et al. 2014). These include marker gene amplicon sequencing (Caporaso et al. 2012), metagenomic and transcriptomic sequencing (Quince et al. 2017, Stewart et al. 2018), multivariate statistical analysis (Buttigieg and Ramette 2014), network analysis (Weiss et al. 2016), and bioinformatics tools (Douglas et al. 2020, Langille et al. 2013), etc. The massive amount of data and knowledge accumulated over the past decades has motivated the development of modeling approaches for a predictive understanding of the functions and interactions within communities (Larsen et al. 2012a, Lopatkin and Collins 2020). Two distinct approaches have been used to model microbial communities: mechanistic modeling and data-driven modeling (Widder et al. 2016, Yao et al. 2022).

Mechanistic modeling can provide insight into the interactions between microbial populations by simulating their growth kinetics (Kumar et al. 2019, Song et al. 2014). Originally used to describe the growth of a single organism on a given substrate (Kovárová-Kovar and Egli 1998, Monod 1942), microbial growth kinetics has been extended to simulate the overall growth of a guild (i.e., a group of populations collectively responsible for a function) on a class of substrates (e.g., organic carbon) (Veshareh and Nick 2021). For example, communities in activated sludge system (an engineered system widely used for wastewater treatment) are modeled by simulating the growth kinetics of four guilds: organic-degrading heterotrophs, ammonia-oxidizing autotrophs, denitrifying bacteria, and phosphate-accumulating organisms (Gujer et al. 1995, Gujer et al. 1999, Henze et al. 1987, Henze et al. 1999). This extension has made mechanistic modeling a robust approach for understanding microbial ecology at the guild level (Batstone et al. 2002, Bouskill et al. 2012, Jin and Roden 2011, Wieder et al. 2015, Wieder et al. 2013). However, the growth kinetics of guilds provide little information about the functions and interactions of individual populations observed in the community. Inferring the functions and interactions of individual populations by simulating their growth kinetics remains challenging due to the lack of knowledge about their growth behavior (Ansari et al. 2021, Rinke et al. 2013).

Data-driven modeling can be used to learn the functions and interactions of microbial populations from their abundance (Kumar et al. 2019, Larsen et al. 2015). This is achieved by capturing the latent relationship between populations and their environments using statistical, probabilistic, or machine learning methods (Ghannam and Techtmann 2021, Larsen et al. 2012a, Mowbray et al. 2021). For example, Bayesian networks, a probabilistic graphical model that can reveal causal relationships between variables via directed acyclic graphs (Uusitalo 2007), have been built to infer microbial interactions and their effects on community structure (Lax et al. 2014, Metcalf et al. 2016, Yuan et al. 2017). Despite its potential, the application of data-driven modeling in microbial ecology is limited by the challenges of data collecting and processing. Because population abundance is difficult and expensive to collect, data-driven models are typically built with small datasets of fewer than a hundred samples (Kuang et al. 2016, Larsen et al. 2012b, Lesnik et al. 2020, Lesnik and Liu 2017, Staley et al. 2014, Yuan et al. 2017). Moreover, population abundance is not adequately processed to capture the variability of microbial functions and interactions at different temporal scales (Ruan et al. 2006, Xia et al. 2011). These challenges can severely compromise the robustness of data-driven inference (Althnian et al. 2021, Ghannam and Techtmann 2021).

Here, a new modeling approach is proposed to leverage mechanistic and data-driven modeling to learn the functions and interactions of individual populations from the growth kinetics of guilds. The mathematical foundation of the proposed approach is revealed by the following derivation. In mechanistic modeling, the total abundance of a guild (*X*) is simulated as first-order kinetics (without considering flow for simplicity):

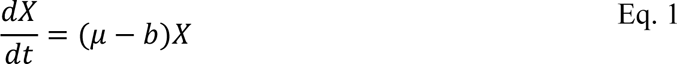

where *µ* and *b* are the growth and decay rates, respectively. The integration of Eq. 1 yields the explicit expression of the specific growth rate of the guild within any given time span (Δ*t*):

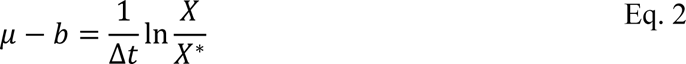

where *X*\* is the historical abundance the guild. The specific growth rate *µ* can be further expressed as a function of substrate concentration using the Monod equation (Monod 1942, 1949):

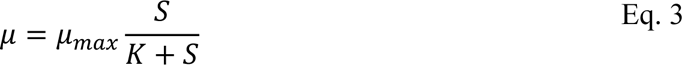

where *µ_max_* is the maximum specific growth rate, *K* is the Monod constant, and *S* is the substrate concentration. Assuming that the guild is composed of *n* populations, the abundance of the guild is the sum of the abundances of all individual populations (*X_i_*) associated with the guild:

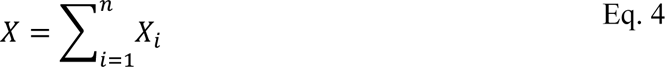

Combining Eqs. 2-4 yields:

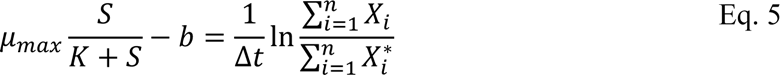

where 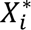 is the historical abundance of the *i*-th population within the guild. Eq. 5 indicates that, within any given time span Δ*t*, the growth kinetics of a guild (numerically represented by its microbial kinetic parameters) are related to the dynamics of individual populations (numerically represented by 1/Δ*t* · ln *X*_*i*_/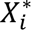). Such an intrinsic connection can be learned using data-driven approaches to infer if a population contributes to the function of the guild and how it interacts with other populations within the guild.

In this study, the proposed modeling approach was implemented based on activated sludge systems. This engineered system has been applied worldwide to treat wastewater, and the Activated Sludge Model No.1 (ASM1) is a well-established mechanistic model that can be readily used to derive the microbial kinetic parameters of guilds (Henze et al. 2000). Meanwhile, activated sludge communities have been extensively studied, and the functional populations in the communities are well curated (McIlroy et al. 2017, Nierychlo et al. 2020, Saunders et al. 2016, Wu et al. 2019). Therefore, activated sludge systems represent an ideal level of complexity for linking microbial growth kinetics to microbial population dynamics. To implement the modeling approach, time series data from four full-scale activated sludge systems were retrieved from the literature. A data transformation method was developed to augment the data and capture the temporal variability of microbial functions and interactions as required by Eq. 5. After data transformation and processing, Bayesian networks were trained to infer functions, and inferences were validated with the

Microbial Database for Activated Sludge (MiDAS) (Dueholm et al. 2022). Interactions were further inferred using topological data analysis. Inferences were also validated indirectly by predicting community structure. Implemented based on activated sludge systems, the modeling approach is expected to be broadly applicable to other natural and engineered ecosystems to provide predictive insight into the functions and interactions within communities.

## 2. Materials and Methods

### 2.1 Data collection and preprocessing

A total of 466 samples were retrieved from three studies to develop the modeling approach (Jiang et al. 2018, Peces et al. 2022, Sun et al. 2021). These studies were selected because they comprehensively reported the environmental factors of four full-scale activated sludge systems in the form of time series (Table 1) and properly deposited sequencing data of 16s rRNA gene amplicon in the National Center for Biotechnology Information database. The diversity of the samples was analyzed using principal coordinate analysis (PCoA) based on Bray-Curtis distance. Sequencing data were retrieved from the database and preprocessed as previously described (Cheng et al. 2021b). Briefly, the QIIME2 pipeline was loaded with DADA2 for denoising the reads (Bolyen et al. 2019). The Silva database (LTPs132_SSU.arb for 16s rDNA, updated in June 2018) was used for sequence classification (Quast et al. 2013). The raw sequences were classified using A classifier that was trained on OTU sequences at 97% identity from the database. The communities in the four activated sludge systems were compared with those from other activated sludge systems using Bray-Curtis distance-based PCoA (Gómez-Acata et al. 2017, Ju and Zhang 2015a, Liu et al. 2017, Ouyang et al. 2016, Saunders et al. 2016, Yuan et al. 2019b, Zhang et al. 2017).

**Table 1.** Overview of the four full-scale activated sludge systems.

|  | Hong Kong | Beijing-1 | Beijing-2 | Aalborg |
| --- | --- | --- | --- | --- |
| **Location** | (22°N, 114°E) | (40°N, 117°E) | (40°N, 110°E) | (57°N, 10°E) |
| **Q (m<sup>3</sup>/day)** | 2.6×10<sup>5</sup> | 1.0×10<sup>6</sup> | 1.0×10<sup>5</sup> | 1.4×10<sup>5</sup> |
| **Sampling Period** | 2013 - 2014 | 2015 - 2016 | 2015 - 2016 | 2017 - 2020 |
| **Number of samples** | 260 | 42 | 42 | 122 |
| **BOD (mg/L)** | 110 - 360 | 180 - 400 | 230 - 1300 | 60 - 430 |
| **Temperature (°C)** | 7 - 31 | 15 - 27 | 15 - 29 | 8 - 22 |
| **SRT (day)** | 9 - 19 | ~10 | ~17 | 10 - 28 |
| **MLVSS (mg/L)** | 1400 - 3300 | 1500 - 3300 | 3500 - 9900 | 1900 - 3900 |

Q: treatment capacity; BOD: biochemical oxygen demand; SRT: sludge retention time; MLVSS: mixed-liquid volatile suspended solid.

Core populations were selected at the family level based on average relative abundance and occurrence frequency across all 466 samples (Ling et al. 2016, Yuan et al. 2019b). More details about core population selection can be found in the Supporting Information (SI) Methods. Redundancy analysis (RDA) was performed on relative abundance and environmental factors. Absolute abundances of core populations were calculated by multiplying relative abundances by volatile suspended solids, a wastewater quality parameter commonly used to represent biomass concentration (Rittmann and McCarty 2012, Saunders et al. 2016).

### 2.2 Data transformation

The 466 samples were randomly divided into training (380 samples) and test (86 samples) sets. Both sets were transformed using the method outlined in Figure 1. First, the time span (Δ*t*) was varied from 7 to 14 days based on the assumption that they could adequately reflect the dynamics of individual populations and the growth kinetics of guilds (Weissman et al. 2021). At each time span, historical and current samples were drawn from the raw datasets to form transformed samples. Taking Δ*t* = 7 d as an example (Figure 1), samples at *t* = 1 (historical) and *t* = 7 (present) were combined to form the first transformed sample, samples at *t* = 2 (historical) and *t* = 8 (present) were combined to form the second transformed sample, and so on. Transformation is repeated for the four activated sludge systems and eight time spans, followed by combining all transformed samples. The 380 raw samples in the training set and 86 samples in the test set were converted into 936 and 134 transformed samples, respectively.

**Figure 1.**
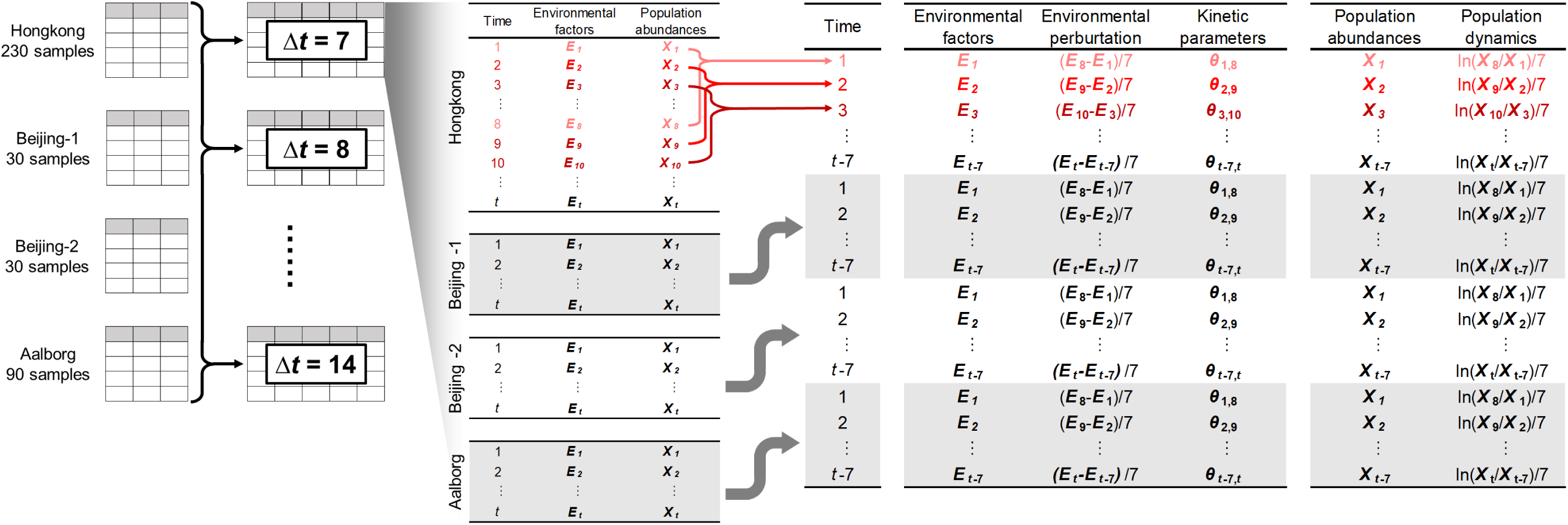
Schematic of applying the data transformation method to the 380 samples in the training set.

Following the transformation, several new features were derived to capture the variability of microbial functions and interactions (Figure 1). First, environmental perturbations, key drivers of population dynamics (Griffin and Wells 2016, Vuono et al. 2014, Yuan et al. 2019a), were quantified as the change in environmental factors normalized to the time span. Second, population dynamics was quantified as the natural logarithm of present versus historical absolute abundances normalized to the time span (SI Method). RDA was performed on environmental perturbation and population dynamics. Finally, microbial kinetic parameters of guilds were derived by inputting the environmental factors within a time span into the ASM1 (SI Methods) (Cheng et al. 2024, Cheng et al. 2021a). The ASM1 was chosen over other activated sludge models because the selected studies only reported data related to heterotrophic organic removal and autotrophic ammonium oxidation. The derived parameters included the maximum specific growth rate *µ_max_*, Monod constant *K*, biomass yield *Y*, and decay rate *b* of heterotrophs and autotrophs.

### 2.3 Model construction

Bayesian networks were built to learn the intrinsic connection between growth kinetics and population dynamics (Eq. 5). The first step of network construction was to scale the values of variables, including time span, historical environmental factors, historical population abundances, environmental perturbation, microbial kinetic parameters and population dynamics, to between 0 and 1 (Bishop 1995):

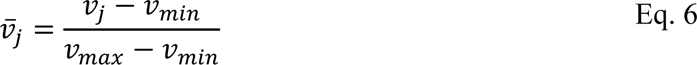

where *v̅_j_* and *v_j_* are the normalized and actual values of variable *v* in sample *j*, respectively; *v_min_* and *v_max_* are the minimum and maximum values of variable *v*, respectively. Next, network structure learning was performed using the R package “bnlearn” with a hill-climbing algorithm and the following assumptions (Scutari et al. 2014): 1) each variable is normally distributed, and variables are multivariate normal; 2) stochastic dependencies are assumed to be linear; 3) time span, historical environmental factors, historical population abundances and environmental perturbation are root parent nodes and cannot depend on any other nodes; 4) for any given population, its dynamics strictly depends on its historical abundance. A final Bayesian network was obtained by averaging 100 bootstrap replicates. Arc strengths representing the frequency of presence in all networks were exported with a threshold calculated by a built-in function in “bnlearn”. Based on the Bayesian inferences, topological data analysis was performed to further capture the microbial interactions within the heterotrophic and autotrophic guilds (SI Methods).

### 2.4 Inference validation

The inferred functions (indicated by the dependencies between kinetic parameters and population abundance/dynamics) from the Bayesian network were validated by searching individual populations in the MiDAS. Within each family, the three most abundant genera with a relative abundance reaching 0.1% were mapped to their corresponding functions in the MiDAS, with particular emphasis on their contribution to heterotrophic organic degradation and autotrophic ammonia oxidation (Dueholm et al. 2022).

As an additional validation step, artificial neural networks (ANNs) were trained to predict population abundance and community structure. To prepare for ANN training, the root parent nodes of population dynamics were identified using a depth-first search algorithm (Heineman et al. 2008). ANNs were then trained with root parent nodes and population dynamics as inputs and outputs, respectively. The hyperparameters of the ANNs were optimized via grid search using the R package Neuralnet (Wright 2019). To perform the optimization, the transformed training set (936 transformed samples) was first divided into 13 subsets (72 transformed samples in each subset). A series of ANNs were then trained with the combinations of different number of hidden layers (2 - 5) and number of nodes (12, 18, 24). The prediction performance of each ANN was evaluated with relative root mean square error (RMSE) based on 13-fold cross-validation (Anguita et al. 2012, Roberts et al. 2017):

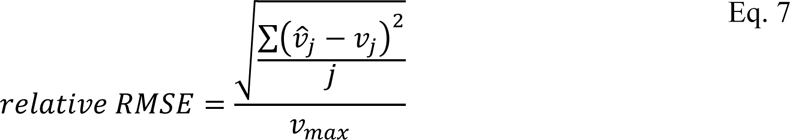

where *v̂_j_* is the predicted value of variable *v* in sample *j*. After hyperparameter optimization, the ANNs with optimal configurations were tested on 134 transformed samples. The ANN outputs (i.e., population dynamics) were converted back to population abundance using historical abundance and time span. The Bray-Curtis similarity between the predicted and observed communities was then calculated.

Two control models were built to compare the prediction performance. The first one was built following previous studies without transforming the data or incorporating microbial growth kinetics (Kuang et al. 2016, Larsen et al. 2012b). A second one is a null model constructed with the mean values of the variables (Harvey et al. 1983).

## 3. Results

### 3.1 Overview of the activated sludge systems

The four activated sludge systems selected for model development were operated under distinct conditions (SI Figure S1). For example, sludge retention time, a key operating factor for microbial community structure (Mansfeldt et al. 2019, Yuan et al. 2019a), varied from 10 days in the Beijing-1 system to 17 days in the Beijing-2 system. The mean temperature and dissolved oxygen were higher in Hong Kong and Beijing than in Aalborg (t-test, *p* < 0.05, SI Figure S1). Moreover, the BOD (600 mg/L) and ammonia (50 mg/L) concentrations in the influent of Beijing-2 were significantly higher than those of Hong Kong (190 and 30 mg/L), Beijing-1 (260 and 45 mg/L), and Aalborg (200 and 30 mg/L) (t-test, *p* < 0.05).

The performance of the systems was also noticeably different (SI Figure S1). Specifically, the effluent BODwas much higher in the two Beijing systems (> 30 mg/L) than in the other two systems (< 5 mg/L), whereas the effluent ammonium showed an opposite trend. On the other hand, the biomass concentration (represented by volatile suspended solids) in Beijing-2 was more than two times higher than in the other systems. As shown by PCoA (Figure 2), the two Beijing systems were significantly different despite their geographic affinity. Although samples from Beijing-1 overlapped with those from Hong Kong and Aalborg, they shifted toward different directions during the seasonal variation (SI Figure S2). Overall, the results show a high diversity of the samples, which is desirable for building robust models.

**Figure 2.**
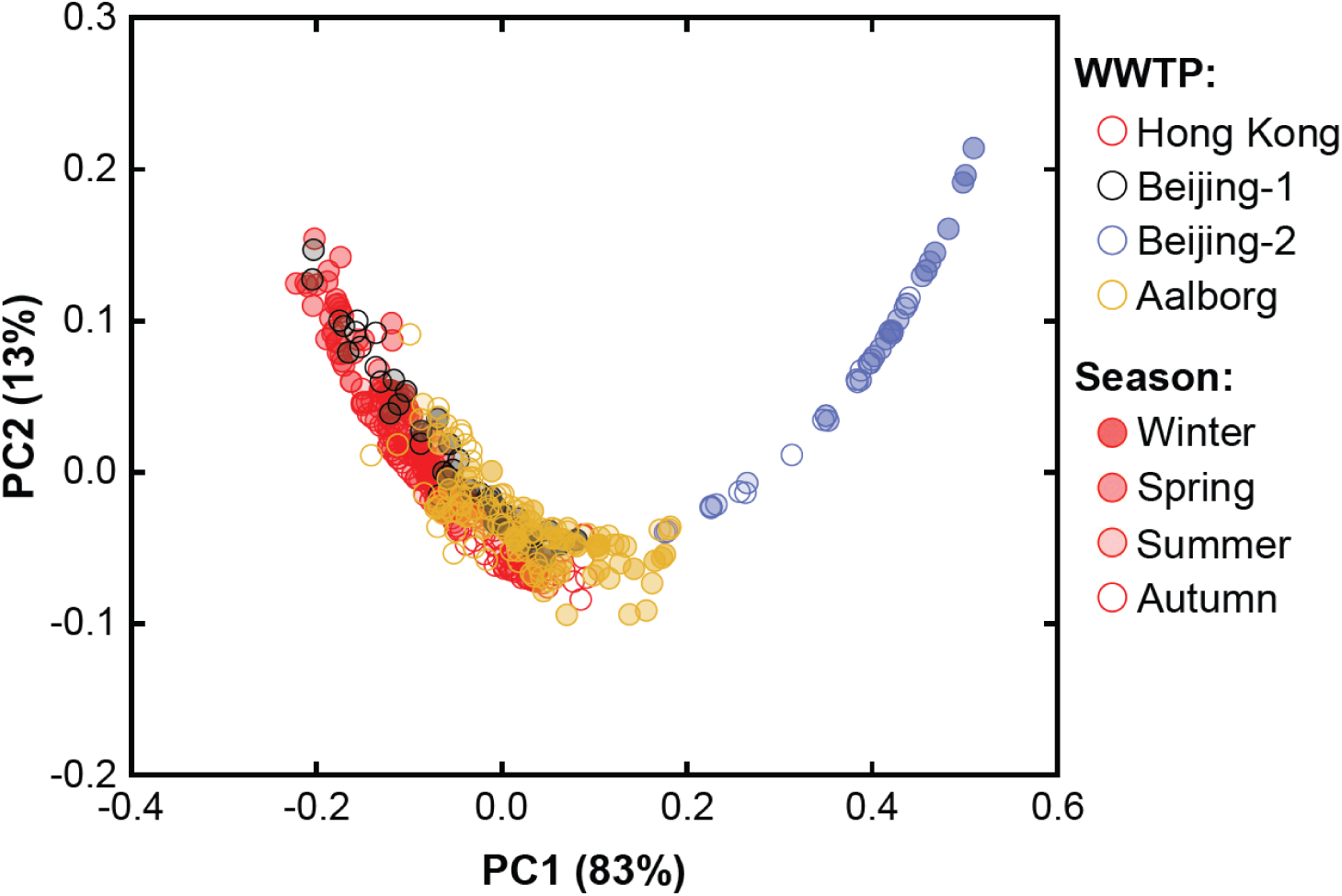
Bray-Curis distance-based PCoA of the environmental factors of the four activated sludge systems.

### 3.2 Overview of the activated sludge communities

The communities in the four systems showed location-dependent differentiation (SI Figure S3). The communities in the two Beijing systems, despite their difference in environmental factors, clustered together and were separate from those in Hong Kong and Aalborg. Similar results were observed in a comprehensive survey of global activated sludge microbiomes (Wu et al. 2019). The communities in Hong Kong and Aalborg showed clear shifts, which could be driven by deterministic factors such as sludge retention time, temperature, and influent concentrations. As shown in SI Figure S1, sludge retention time fluctuated more dramatically in Hong Kong and Aalborg than in Beijing. This environmental factor has been shown to exert strong selective pressure and drive deterministic community assembly (Mansfeldt et al. 2019, Vuono et al. 2014). Although the communities in the three sites formed individual clusters, they belonged to an activated sludge microbiome formed by eleven systems in China, Mexico, the U.S., and Denmark, demonstrating the representativeness of the communities being modeled.

A total of 42 core populations were selected at the family level based on the following criteria: average abundance > 0.5% and occurrence > 35% across all 466 samples (SI Methods and Table S1). They accounted for an average of 70% of the total relative abundance and included common activated sludge populations such as Comamonadaceae, Rhodobacteraceae, Saprospiraceae, Nitrospiraceae, Nitrosomonadaceae, and Rhodocyclaceae (Figure 3). As revealed by RDA, Comamonadaceae, Rhodobacteraceae and Saprospiraceae were influenced by organic carbon (i.e., BOD) and dissolved oxygen (SI Figure S4). In comparison, Nitrospiraceae, Nitrosomonadaceae and Rhodocyclaceae showed high sensitivity to ammonia and temperature (SI Figure S4). Members of these families were frequently found abundant in activated sludge communities and were known to contribute to organic carbon degradation, ammonia/nitrite oxidation, and nitrate reduction (Dueholm et al. 2022). The core also included less well characterized populations (e.g., Bradyrhizobiaceae, Sphingomonadaceae, and Intrasporangiaceae), as well as unknown taxa due to the lack of their representative sequences in the *Silva* database (updated in June 2018).

**Figure 3.**
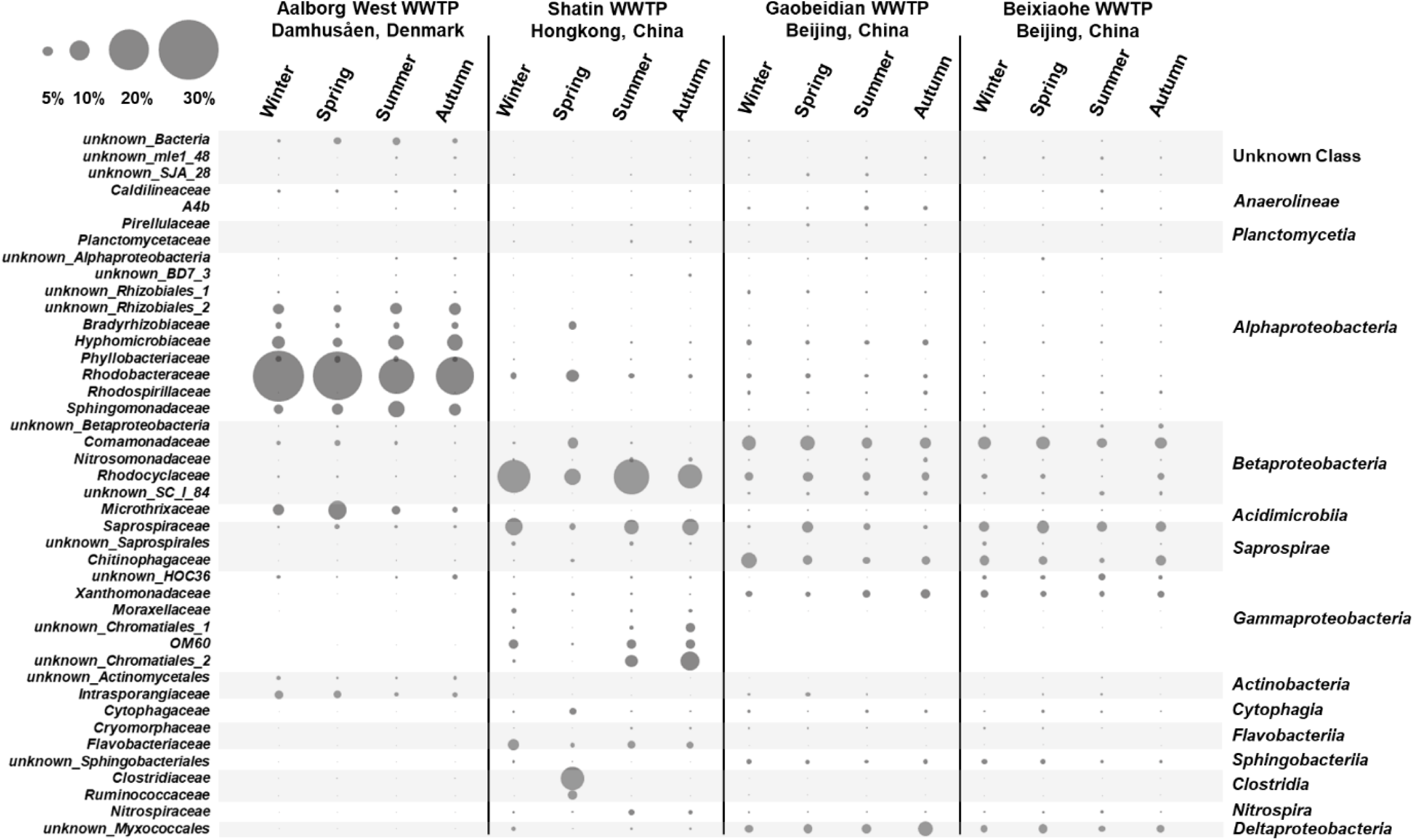
Relative abundance of the 42 core populations selected at the family level in the four activated sludge systems.

### 3.3 Dynamics of the core populations

Population dynamics was expected to aggregate the periodic fluctuation in microbial growth and reflect the stability of the populations (SI Methods). As shown in Figure 4, the majority of the core populations exhibited high stability with the dynamics ranging between ±0.05 d^-1^. Comamonadaceae, Rhodobacteraceae and Rhodocyclaceae were particularly stable, as evidenced by the close-to-zero dynamics (−0.015 d^-1^) and narrow 95% confidence intervals (< 0.015 d^-1^) across all 936 transformed samples. These indicated their resilience to environment perturbation and explained their ubiquity in activated sludge systems. Fluctuation was observed for Saprospiraceae (confidence interval >0.02), suggesting their weaker resilience to environmental perturbation. For nitrifying bacteria, Nitrospiraceae appeared to be more stable (−0.03 d^-1^, 95% confidence interval 0.03) than Nitrosomonadaceae (−0.07 d^-1^, 95% confidence interval 0.04) (t-test, *p* < 0.05).

**Figure 4.**
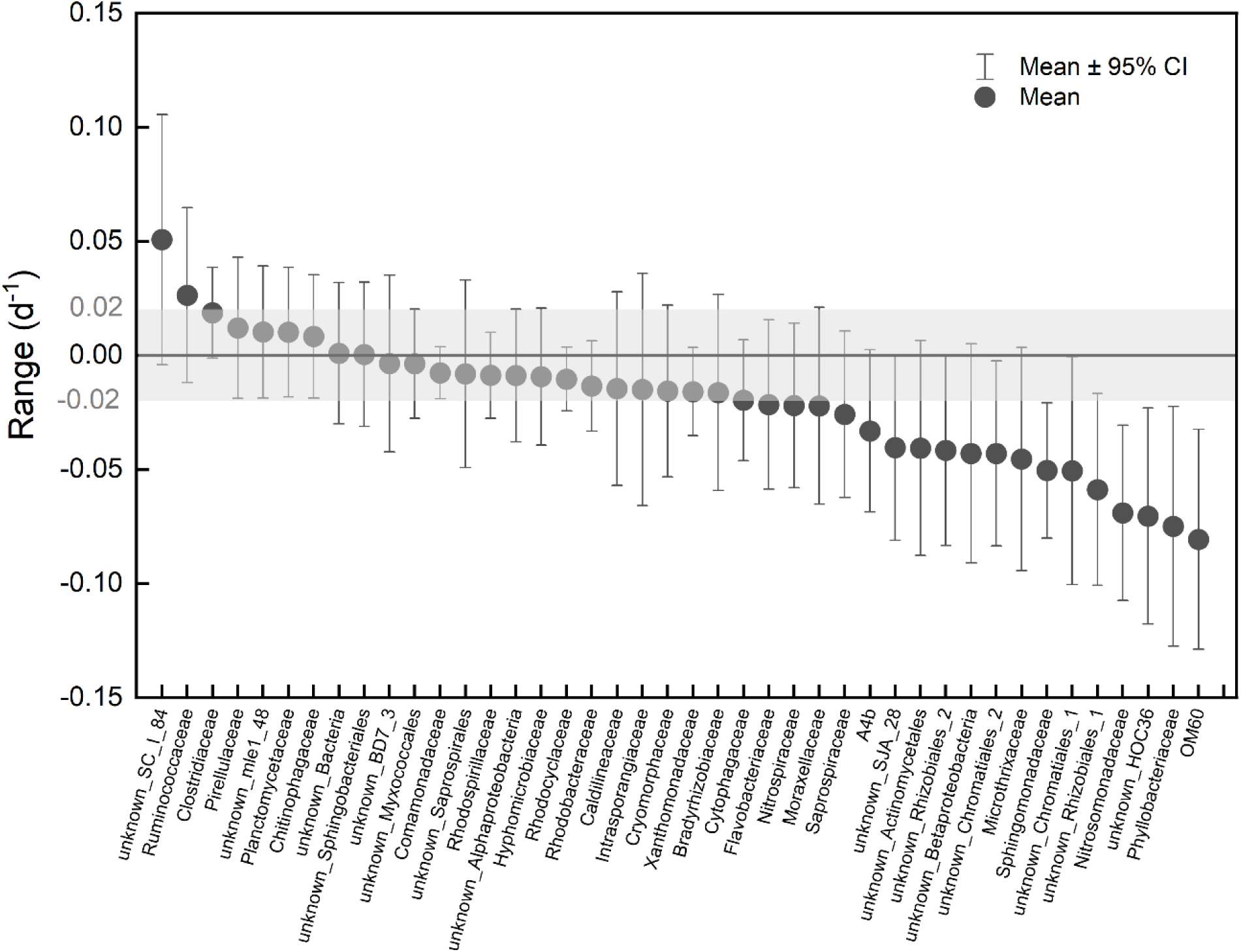
Overall dynamics of the 42 core populations across eight time spans and 936 transformed samples.

RDA was performed to understand the relationship between population dynamics and environmental perturbation (SI Figure S5). Most of the core populations were found in the center of the RDA plot, meaning that they were not influenced by the change in any particular environmental factor. For example, Comamonadaceae, Rhodobacteraceae and Rhodocyclaceae were very close to the origin, highlighting their resilience to perturbation. Saprospiraceae was slightly affected by the changes in sludge retention time and effluent ammonia, consistent with its dynamics (Figure 4). Similarly, Nitrospiraceae was closer to the origin, whereas Nitrosomonadaceae was more positively associated with sludge retention time and negatively associated with temperature. Populations with the high dynamics (e.g., unknown SC_I_84, OM60, Phyllobacteriaceae) were found in the outermost areas in the RDA plot and were strongly affected by environmental perturbation.

### 3.4 Inference of functions and interactions

Bayesian networks were built through data-driven structure learning to identify associations between growth kinetics and population dynamics. The subnetwork centered on heterotrophic growth kinetics consisted of typical heterotrophs such as Comamonadaceae, Rhodobacteraceae, and Saprospiraceae (Figure 5A). Comamonadaceae were directed to the Monod constant of aerobic heterotrophs (*K_oh_*) with an arc strength of 0.61 (SI Table S2). According to the MiDAS, the dominant genera in this family were all identified as aerobic heterotrophs and could grow on complex substrates such as sugars and proteins (SI Figure S7). Rhodobacteraceae served as an alternative aerobic heterotroph, as evidenced by its strong dependence on the maximum specific growth rate of heterotrophs (*u_h_*, arc strength 0.75). The genera of Rhodobacteraceae were abundant and could also degrade different substrates (SI Figure S7). Saprospiraceae was associated with the biomass yield of aerobic heterotrophs (*Y_h_*) with an arc strength of 0.69 (SI Table S2). This family was dominated by an unknown genus (average relative abundance 4.9%) that could not be mapped to the MiDAS. One of its potential functions is to prey on live cells and/or convert dead cells into simpler organic compounds (Seguel Suazo et al. 2024).

**Figure 5.**
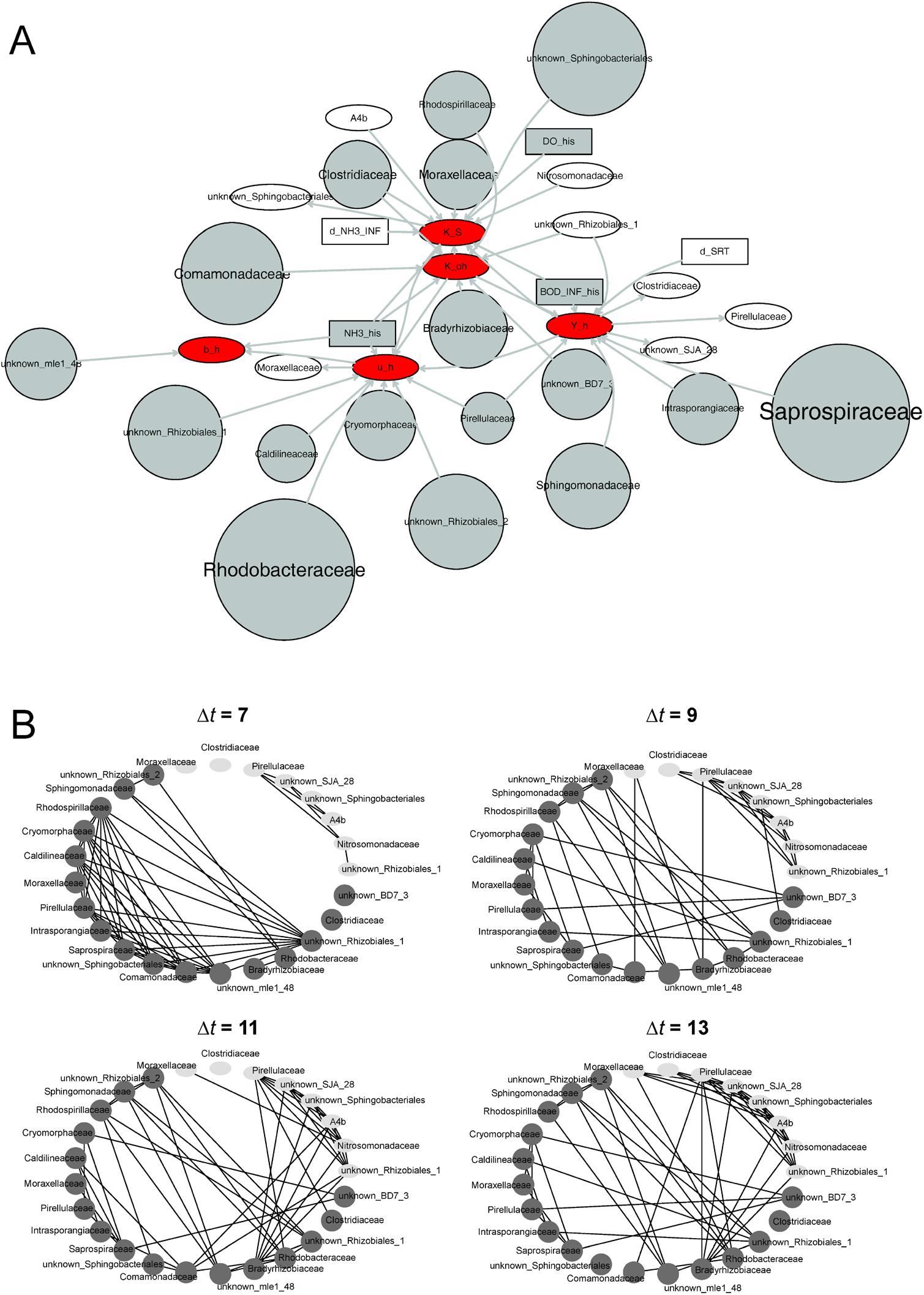
(A) Bayesian network centered on heterotrophic growth kinetics. Red oval indicates microbial kinetic parameters, white oval indicates population dynamics, grey circle indicates historical population abundance, white box indicates environmental perturbation, and grey box indicates historical environmental factors. (B) Topological data analysis at Δt = 7, 9, 11, and 13.

Topological data analysis was performed to further learn the interactions within populations related to heterotrophic growth kinetics (Figure 5B and SI Figure S9). The results showed a time-dependent relationship between historical abundance and population dynamics. Specifically, the average number of arcs between the two categories increased from less than three at short time spans (Δt < 10 days) to more than seven at long time spans (Δt > 11 days). Bradyrhizobiaceae was identified as a keystone population as its historical abundance was linked to the dynamics of multiple populations including Pirellulaceae at different time spans. Bradyrhizobiaceae and Pirellulaceae have been reported to coexist in various environments including marine sediments and soils (Liu et al. 2018, Walters et al. 2018). According to the MiDAS, the dominant genus of Bradyrhizobiaceae, *Bradyrhizobium*, can perform denitrification (SI Figure S7). Within the interactions learned from historical abundance, consistent patterns were observed. For example, Rhodobacteraceae was associated with Sphingomonadaceae across all eight time spans, and their interactions at short time spans were among the strongest, as revealed by the adjacency matrices (i.e., the inverse of the Euclidean distance, SI Table S3). Similarly, the interaction between Saprospiraceae and Cryomorphaceae were consistently observed in the top ten adjacency list.

In the subnetwork centered on autotrophic growth kinetics (Figure S6), Nitrospiraceae and Nitrosomonadaceae were found to point to the decay and growth rates of autotrophs with an arc strength of 0.64 and 0.59, respectively (SI Table S4). The genus *Nitrospira* of Nitrospiraceae is known to be involved in ammonia and nitrite oxidation (SI Figure S8). In the family Nitrosomonadaceae, an unknown genus contributed 99% of the abundance but could not be found in the MiDAS. This genus is likely an ammonium/nitrite oxidizer but warrants further investigation. Rhodocyclaceae as the most abundant population was associated with the Monod constant of ammonia oxidation (*Kn*) with an arc strength of 0.60 (SI Table S4). According to the MiDAS, the genera in this family can suppress ammonia- and nitrite-oxidizing bacteria (e.g., *Candidatus* Accumulibacter, SI Figure S8). Therefore, the presence of Rhodocyclaceae is expected to exert negative effects on nitrification. In addition, families such as Sphingomonadaceae, Intrasporangiaceae, and Caldilineaceae were associated with both the microbial kinetic parameters of heterotrophs (Figure 5A) and autotroph (Figure S6). Similar to Rhodocyclaceae, these populations are listed as aerobic heterotrophs in the MiDAS (SI Figure S8) and potentially inhibit nitrification.

Topological data analysis of the autotroph-related populations revealed a clear independence between historical abundance and population dynamics (SI Figure S10). The interactions within the autotrophic guild were also simpler than those between the heterotroph-related populations as evidenced by the greater sparsity of the arcs. Moreover, the interactions appeared to be more consistent across time spans. For example, when interactions were inferred from historical abundance, Nitrospiraceae was always associated with Rhodocyclaceae and an unknown family of the order Chromatiales, while Rhodocyclaceae interacted consistently with Moraxellaceae and OM60. In terms of the dynamics-inferred associations, Nitrosomonadaceae and Caldilineaceae actively interacted at all time spans with the adjacency among the highest at most time spans (SI Table S5). Meanwhile, Nitrosomonadaceae, Intrasporangiaceae, and OM60 formed a stable relationship throughout the eight time spans (SI Figure S10). Their adjacencies were ranked in the top ten at time spans 7 and 9-14 (SI Table S5).

### 3.5 Validation of the inferences

The inferred functions and interactions were indirectly validated by predicting absolute abundance and community structure, with the expectation that adequate inferences should lead to accurate predictions. After optimization using a grid search method (SI Methods), ANNs were tested on 134 transformed samples to predict absolute abundance and community structure. A total of 5628 predictions (134 samples × 42 populations) were obtained (SI Figure S11). The fitted slope of 0.51 between observed and predicted abundances indicated that the model tended to overestimate population abundance. This was because a large number of populations were absent (i.e., zero abundance) in the test set but were predicted to be present with certain abundances. As a result, the R^2^ between observed and predicted abundances was only 0.36. When the absent populations were excluded from the test set, the prediction was significantly more accurate with the slope and R^2^ increasing to 0.75 and 0.56, respectively. In comparison, the ANNs trained without microbial kinetic parameters as inputs yielded more server overestimation (slope 0.47) and less accurate predictions (R^2^ 0.50) even when the absent populations were excluded. The difficulty in predicting the abundance of individual populations, especially those with low to zero abundances, could be attributed to the incompleteness of the dataset (e.g., lack of environmental factors such as nitrate and phosphate concentrations and their perturbation), as well as the inherently stochastic processes of microbial community assembly (Langille et al. 2013, Zhou and Ning 2017).

The predicted and observed community structures were compared using Bray-Curtis similarity. The communities predicted with microbial kinetic parameters as inputs shared a similarity of 0.70 ± 0.19 with the observed communities (Figure 6). This was comparable to the results obtained in previous studies (e.g., 0.65 for order-level communities in acid mine drainage and 0.72 for genus-level communities in bioelectrochemical systems) (Cheng et al. 2021a, Langille et al. 2013). In comparison, prediction without microbial kinetic parameters yielded a lower similarity of 0.66 ± 0.18 (t-test, *p* < 0.05). Given that over half of the raw samples were from Hong Kong, additional predictions were performed for the Hong Kong samples only. The similarity increased significantly to 0.84 ± 0.07 (with parameters) and 0.70 ± 0.20 (without parameters). The results suggest slight overfitting caused by inherent bias in the training set, as well as the benefit of including microbial kinetic parameters to mitigate such bias. To further understand the potential of the model, ANNs were trained and tested with relative abundance as input. Compared to those trained with absolute abundance, the similarity was improved to 0.74 ± 0.17 (SI Figure S12, t-test, *p* < 0.05). The improvement could be due to the elimination of the error introduced by biomass concentration. The control model (built following a previous study (Larsen et al. 2012b)) and the null model (built using average abundances) produced much less accurate predictions with a similarity of 0.60 ± 0.20 and 0.53 ± 0.19, respectively (t-test, *p* < 0.05).

**Figure 6.**
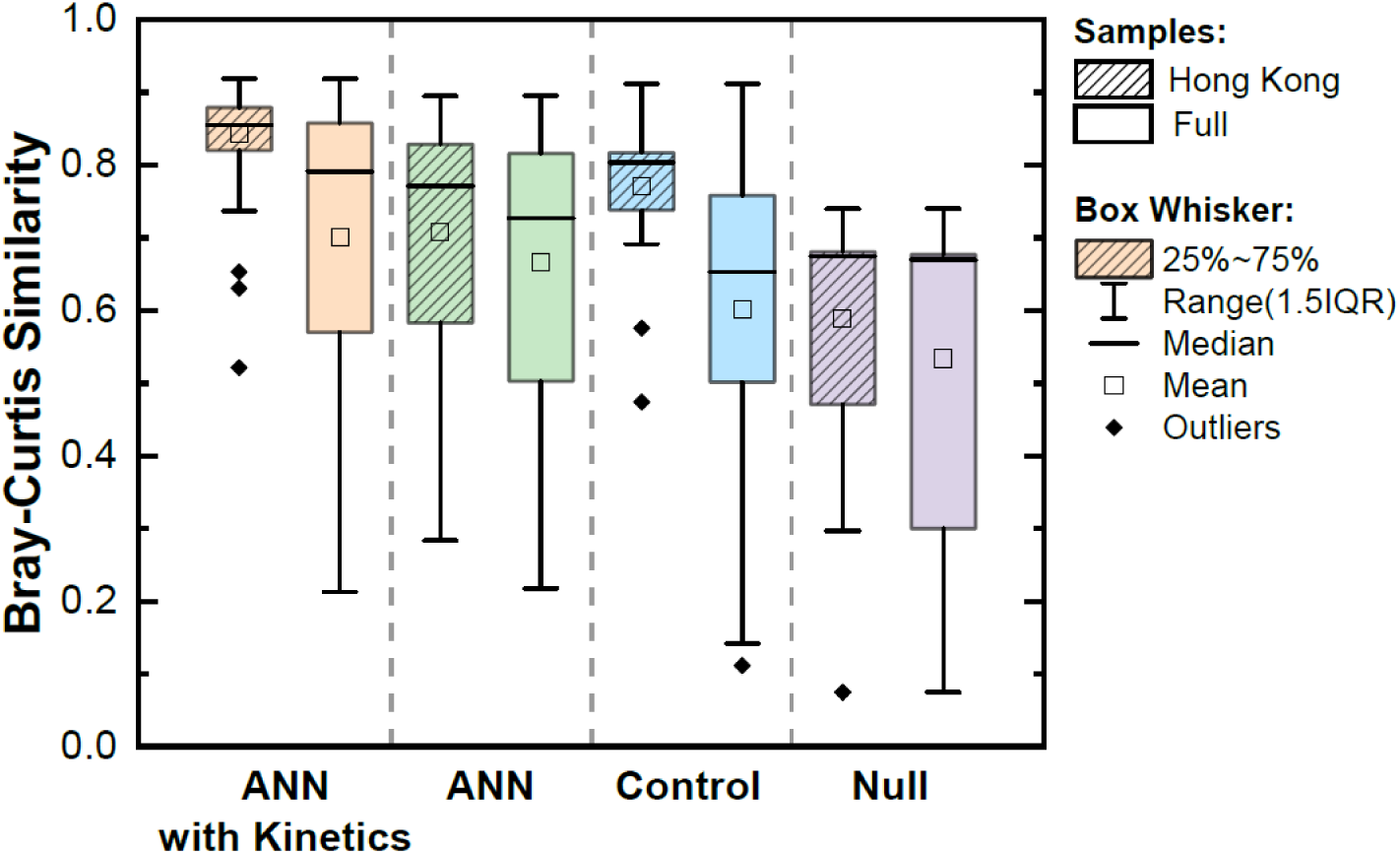
Bray-Curtis similarity between predicted and observed communities.

## 4. Discussion

There is a growing and extensive interest in building a predictive understanding of microbial ecology by modeling microbial communities (Ghannam and Techtmann 2021, Kumar et al. 2019, Larsen et al. 2012a, Lopatkin and Collins 2020). Conventional mechanistic models can predict the functions and interactions of guilds but not the actual microbial populations observed in the community due to the challenges of simulating the growth kinetics of individual populations (Bouskill et al. 2012, Song et al. 2014). Emerging data-driven models can infer the functions and interactions within the community, but the robustness of the inference can be compromised by limited data availability and lack of temporal variation (Metcalf et al. 2016, Yao et al. 2022).

The present study aimed to address these limitations by integrating mechanistic and data-driven modeling based on the intrinsic connection between microbial growth kinetics and microbial population dynamics (Eq. 5). To implement this novel modeling approach, a comprehensive literature review was conducted, and 466 raw samples from four full-scale activated sludge systems in Hong Kong, Beijing, and Aalborg were retrieved (Table 1) (Jiang et al. 2018, Peces et al. 2022, Sun et al. 2021). The environmental factors and microbial communities were highly diverse (Figures 2 and 3), demonstrating the representativeness of the selected samples for implementing the modeling approach. Using the data transformation method developed in this study (Figure 1), samples from different activated sludge systems were transformed into the same structure and combined, and the size of the training set was increased by three times.

In addition to data augmentation, the transformation enabled quantification of population dynamics and environmental perturbation. The overall dynamics of a core population that is universally abundant in different environments is expected to be close to zero (SI Methods). Consistent with this assumption, 34 of the 42 core populations exhibited low dynamics within ±0.05 d^-1^ (Figure 4) and were resilient to environmental perturbation (SI Figure S4). Although the calculation of population dynamics differs from a previously developed mass balance-based method that calculates net growth rates (Kim et al. 2020, Peces et al. 2022, Saunders et al. 2016), the two methods are complementary in terms of identifying consistently active populations. The remaining eight populations were highly dynamic and strongly affected by environmental perturbation (SI Figure S6). Notably, unknown SC_I_84 with the most positive dynamics showed the strongest positive association with temperature perturbation and negative association with sludge retention time perturbation, whereas the populations with the most negative dynamics (Nitrosomonadaceae, Phyllobacteriaceae, and OM60) showed the exact opposite. The results imply the replacement of the latter by the former and highlight the potential of the data transformation method for holistic analysis of microbiomes from different systems, which can lead to new insights into microbial ecology (Wu et al. 2019).

Learned from the transformed data, the Bayesian network and topological associations served as exploratory tools for identifying keystone populations and inferring their roles within the community. For example, the family Intrasporangiaceae was found to be linked to the parameters of both heterotrophs and autotrophs (SI Figure S5). Dominated by the putative phosphate-accumulating organism *Tetrasphaera* (Kristiansen et al. 2013), Intrasporangiaceae was consistently associated with Nitrosomonadaceae in topological data analysis (SI Figure S10). Together with their dynamics and the MiDAS, it is reasonable to speculate that Intrasporangiaceae and Nitrosomonadaceae have an agonistic relationship. Such inferences can be combined with downstream ANN prediction of population abundance to forecast system performance. For example, high abundance of suppressors such as Intrasporangiaceae and low abundance of nitrifying bacteria would result in excess nitrates and nitrites, which in turn could inhibit various microorganisms depending on pH, temperature, and influent concentrations (Zhou et al. 2011). This inhibition could lead to significant shifts in microbial community structure and consequently impair system performance. Predicted dominance and washout of indicator populations can alert treatment plant personnel and enable predictive control to prevent system failure.

Despite their potential to provide predictive insights into microbial ecology and engineering applications, the data transformation method and modeling approach developed in this study remain to be improved with more data, particularly time series of population abundance collected at high sampling frequencies (e.g., every 1-3 days). Several global surveys have been conducted to understand the microbiomes in activated sludge systems and anaerobic digesters (Mei et al. 2017, Wu et al. 2019), but the data are not reported as time series and cannot be incorporated into this study. The modeling approach also needs to be examined with natural ecosystems, where the growth kinetics of guilds have been simulated mechanistically (Bouskill et al. 2012, Jin and Roden 2011, Wieder et al. 2015, Wieder et al. 2013). Finally, the modeling approach can be improved by proper selection and integration of probabilistic and machine learning methods. In this study, Bayesian networks have two functions: to classify populations based on probabilistic dependencies with guild kinetic parameters, and to reduce the computational cost of downstream topological data analysis and ANN training. Topological data analysis has emerged as a powerful method in machine learning/deep learning for extracting complementary information about the observed objects (Baas et al. 2020, Wasserman 2018, Zomorodian 2012). It is applied for the first time to capture hidden spatial dependencies and genuine associations between microbial populations, and its application in microbial ecology remains to be refined.

## Acknowledgment

This work was supported by the U.S. Department of Agriculture [Award No. 2020-67019-31027].

