## Supplementary material for "Building Predictive Understanding of Microbial Ecology by Bridging Microbial Growth Kinetics and Microbial Population Dynamics": AS Modeling_SI.docx

Intended for ***TBD***

Type of contribution***: Research Article***

* **Corresponding author**

Yuzhou Chen:

Heyang Yuan:

**SI Methods**

**Core population selection**

Core populations were selected based on two criteria: average relative abundance and frequency of occurrence in all 466 samples (Yuan et al. 2019, Cheng et al. 2021). To identify the optimal combination of these criteria, core populations were selected with a series of average relative abundance and occurrence. The average relative abundance ranged from 0.1%, 0.3%, 0.5%, 0.7%, to 1%, and the occurrence ranged from 10%, 30%, 35%, 50%, and 70% (SI Table S1), resulting in a total of 25 combinations. To mitigate the bias resulting from the high weight of samples under specific conditions (e.g., from the same activated sludge systems, with similar sludge retention time, in the same season, etc.), the calculated average relative abundance and occurrence frequency were adjusted based on the number of samples under specific conditions.

**Population dynamics calculation**

The dynamics of a population was quantified as the natural logarithm of present versus historical absolute abundances normalized to time span:

$$Dynamics=\frac{1}{\Delta t}\ln\frac{X}{X^{*}}$$

Eq. S1

The overall dynamics of a population was calculated by averaging its dynamics obtained at eight time spans. Population dynamics can reflect microbial growth, as revealed by the following equations. The mass balance of any given microbial population can be expressed as:

$$V\frac{dX}{dt}=Q_{in}X_{in}-Q_{out}X_{out}-Q_{w}X_{w}+V\mu X$$

Eq. S2

The modeling boundary was set from influent to secondary settling tank effluent. The influent cell concentration is negligible ($X_{in}\approx0$), while the effluent cell concentration is almost removed due to biomass settling ($X_{out}\approx0$). The term $Q_{w}X_{w}$ is the cell wasting rate and depends on the biomass in activated sludge basin. Thus, ${Q_{w}X}_{w}$ can be expressed as the product of $X$ and a concentration factor $c$ (L/d). Eq. S2 can then be reduced to:

$$V\frac{dX}{dt}=-Q_{w}cX+V\mu X$$

Eq. S3

Integration of Eq. S3 yields:

$$\frac{1}{\Delta t}\ln\frac{X}{X^{*}}=\mu-\frac{Q_{w}}{V}c$$

Eq. S4

Therefore, population dynamics is correlated with the net growth rate. When the abundance of population remains stable (i.e., $X_{i}\approx X_{i}^{*}$), its dynamics approaches zero, indicating its resilience against environmental perturbation.

**Kinetic parameter calculation**

After collecting the training dataset, microbial kinetic parameters were calibrated in the activated sludge model No. 1 (ASM1, (Henze et al. 2006)) for subsequent training of the data-driven component. The ASM1 was simplified to simulate a duo-guild interaction (e.g., autotrophs and heterotrophs). The microbial processes were described using the following equation:

$\frac{dS_{S}}{dt}=D(S_{S,in}-S_{S})-\frac{1}{Y_{h}}\mu_{h}(\frac{S_{S}}{S_{S}+K_{S}})(\frac{S_{O}}{S_{O}+K_{O,h}})X_{h}$	Eq. S5

$\frac{dX_{h}}{dt}=\mu_{h}\left( \frac{S_{S}}{S_{S}+K_{S}} \right)\left( \frac{S_{O}}{S_{O}+K_{O,h}} \right)X_{h}-b_{h}X_{h}$	Eq. S6

$\frac{dX_{a}}{dt}=\mu_{a}\left( \frac{S_{NH}}{S_{NH}+K_{NH}} \right)\left( \frac{S_{O}}{S_{O}+K_{O,a}} \right)X_{a}-b_{a}X_{a}$	Eq. S7

$\frac{dS_{O}}{dt}=R_{ar}-\frac{1-Y_{h}}{Y_{h}}\mu_{h}\left( \frac{S_{S}}{S_{S}+K_{S}} \right)\left( \frac{S_{O}}{S_{O}+K_{O,h}} \right)X_{h}-\frac{4.57-Y_{a}}{Y_{a}}\mu_{a}\left( \frac{S_{NH}}{S_{NH}+K_{NH}} \right)\left( \frac{S_{O}}{S_{O}+K_{O,a}} \right)X_{a}$	Eq. S8

$\frac{dS_{NH}}{dt}=D(S_{NH,in}-S_{NH})-f_{b,n}\mu_{h}(\frac{S_{S}}{S_{S}+K_{S}})(\frac{S_{O}}{S_{O}+K_{O,h}})X_{h}-(f_{b,n}+\frac{1}{Y_{a}})\mu_{a}\left( \frac{S_{NH}}{S_{NH}+K_{NH}} \right)\left( \frac{S_{O}}{S_{O}+K_{O,a}} \right)X_{a}$	Eq. S9

Where *S* indicates concentration, and the subscription “*S*”, “*O*”, and “*NH*” represents substrate, oxygen, and ammonia, respectively, and the subscription “*in*” represents influent; $R_{ar}$ is the aeration rate in tank; *X* indicates biomass, and the subscription “*h*” and “*a*” represents heterotrophs and autotrophs; the dilution rate *D* equals $\frac{Q_{in}}{V}$; the coefficient $f_{b,n}$ (0.086 (Gujer et al. 1995, Henze et al. 2006)) is the fraction of ammonia for cell synthesis. The terms in red are the microbial kinetic parameters to be calibrated. These include the maximum specific growth rates *μ*, yield coefficients $Y$, decay rate constants *b*, and half-rate constants $K$ for autotrophs and heterotrophs. The processes related to nitrates were not considered here because the nitrate data were missing in the literature. The discharging rate of waste sludge was also ignored due to the missing data. As a result, this biomass loss was combined into the decay rate constants *b* during the calibration.

**Testing of ANNs**

The 76 samples in the test set were converted into 134 transformed samples. Some of the calibrated values were constant during calibration (e.g., 1 mg/L for the half rate constant K_oa_) because they were the boundary values determined based on previous studies (Henze et al. 2006, Bruce 2020). Due to an addition of 10^-8^ to all the original relative abundance for calculation of population dynamics, the points with observed zero values (i.e., ~900 points around x = 10^-5^) were isolated from the others.

Table S1. Summary of core population selection. The upper table shows the number of populations (at the family level) selected with different criteria. The lower table shows the average total relative abundance (%) of the populations selected.

| **Average  Abundance Frequency** | 0.1 | 0.3 | 0.5 | 0.7 | 1 |
| --- | --- | --- | --- | --- | --- |
| 10 | 134 | 67 | 45 | 35 | 25 |
| 30 | 108 | 63 | 45 | 35 | 25 |
| 35 | 96 | 60 | 42 | 33 | 24 |
| 50 | 63 | 50 | 37 | 29 | 20 |
| 70 | 30 | 30 | 28 | 21 | 16 |
| **Average  Abundance Frequency** | 0.1 | 0.3 | 0.5 | 0.7 | 1 |
| 10 | 92.43 | 80.75 | 72.56 | 66.12 | 57.86 |
| 30 | 87.58 | 79.09 | 72.56 | 66.12 | 57.86 |
| 35 | 83.55 | 76.44 | 69.91 | 64.17 | 56.76 |
| 50 | 70.2 | 67.14 | 62.54 | 57.29 | 49.89 |
| 70 | 53.45 | 53.45 | 52.76 | 48.12 | 44.09 |

Table S2. Strength of arcs that involve heterotrophs’ kinetics in the Bayesian Network.

| **Arcs** | **From** | **To** | **Strength** |
| --- | --- | --- | --- |
| 1 | K_oh | K_S | 0.63 |
| 2 | K_S | u_h | 0.61 |
| 3 | unknown_Rhizobiales_2_his | u_h | 0.62 |
| 4 | K_oh | Y_h | 0.51 |
| 5 | Sphingomonadaceae_his | Y_h | 0.63 |
| 6 | Y_h | u_h | 0.61 |
| 7 | Rhodospirillaceae_his | K_oh | 0.67 |
| 8 | Y_h | K_S | 0.61 |
| 9 | Cryomorphaceae_his | u_h | 0.62 |
| 10 | Caldilineaceae_his | u_h | 0.63 |
| 11 | Moraxellaceae_his | K_S | 0.62 |
| 12 | unknown_Rhizobiales_1 | Y_h | 0.65 |
| 13 | Pirellulaceae_his | u_h | 0.61 |
| 14 | NH3_his | K_oh | 0.61 |
| 15 | Intrasporangiaceae_his | Y_h | 0.64 |
| 16 | Saprospiraceae_his | Y_h | 0.69 |
| 17 | unknown_Sphingobacteriales_his | K_S | 0.63 |
| 18 | NH3_his | K_S | 0.60 |
| 19 | BOD_INF_his | Y_h | 0.65 |
| 20 | K_S | unknown_Sphingobacteriales | 0.59 |
| 21 | Comamonadaceae_his | K_oh | 0.61 |
| 22 | NH3_his | u_h | 0.55 |
| 23 | Time_Span | K_S | 0.59 |
| 24 | d_SRT | Y_h | 0.61 |
| 25 | d_NH3_INF | K_S | 0.65 |
| 26 | Pirellulaceae_his | Y_h | 0.61 |
| 27 | Time_Span | Y_h | 0.59 |
| 28 | u_h | b_h | 0.63 |
| 29 | Y_h | unknown_SJA_28 | 0.58 |
| 30 | Y_h | Pirellulaceae | 0.59 |
| 31 | Y_h | Clostridiaceae | 0.58 |
| 32 | NH3_his | b_h | 0.61 |
| 33 | unknown_mle1_48_his | b_h | 0.62 |
| 34 | Bradyrhizobiaceae_his | K_oh | 0.63 |
| 35 | K_oh | u_h | 0.53 |
| 36 | Rhodobacteraceae_his | u_h | 0.75 |
| 37 | unknown_Rhizobiales_1_his | u_h | 0.60 |
| 38 | Time_Span | K_oh | 0.59 |
| 39 | DO_his | K_S | 0.63 |
| 40 | Clostridiaceae_his | K_S | 0.61 |
| 41 | BOD_INF_his | K_S | 0.63 |
| 42 | u_h | Moraxellaceae | 0.60 |
| 43 | Clostridiaceae_his | K_oh | 0.62 |
| 44 | unknown_Rhizobiales_1 | K_oh | 0.62 |
| 45 | unknown_BD7_3_his | K_oh | 0.60 |
| 46 | Nitrosomonadaceae | K_S | 0.64 |
| 47 | A4b | K_S | 0.63 |

Table S3. Strength of arcs that involve autotrophs’ kinetics in the Bayesian Network.

| **Arcs** | **From** | **To** | **Strength** |
| --- | --- | --- | --- |
| 1 | u_a | K_oa | 0.74 |
| 2 | OM60_his | b_a | 0.63 |
| 3 | Sphingomonadaceae_his | u_a | 0.62 |
| 4 | d_BOD_INF | b_a | 0.76 |
| 5 | Nitrospiraceae_his | b_a | 0.64 |
| 6 | K_n | u_a | 0.59 |
| 7 | d_NH3_INF | b_a | 0.66 |
| 8 | Intrasporangiaceae_his | K_oa | 0.64 |
| 9 | u_a | Pirellulaceae | 0.60 |
| 10 | Ruminococcaceae_his | b_a | 0.62 |
| 11 | unknown_Chromatiales_2_his | K_n | 0.63 |
| 12 | K_n | Y_a | 0.64 |
| 13 | Moraxellaceae_his | K_n | 0.63 |
| 14 | Time_Span | K_n | 0.60 |
| 15 | b_a | Pirellulaceae | 0.59 |
| 16 | unknown_Bacteria_his | K_n | 0.64 |
| 17 | Phyllobacteriaceae_his | u_a | 0.60 |
| 18 | b_a | OM60 | 0.53 |
| 19 | Moraxellaceae_his | b_a | 0.60 |
| 20 | K_n | unknown_SJA_28 | 0.55 |
| 21 | K_n | Pirellulaceae | 0.57 |
| 22 | K_n | b_a | 0.59 |
| 23 | K_oa | unknown_Alphaproteobacteria | 0.58 |
| 24 | DO_his | Y_a | 0.60 |
| 25 | b_a | Moraxellaceae | 0.59 |
| 26 | K_oa | Pirellulaceae | 0.50 |
| 27 | Caldilineaceae | u_a | 0.60 |
| 28 | Nitrosomonadaceae | u_a | 0.59 |
| 29 | A4b_his | K_n | 0.64 |
| 30 | K_n | Intrasporangiaceae | 0.53 |
| 31 | unknown_Rhizobiales_2_his | K_oa | 0.60 |
| 32 | d_NH3_INF | K_n | 0.65 |
| 33 | Rhodocyclaceae_his | K_n | 0.60 |

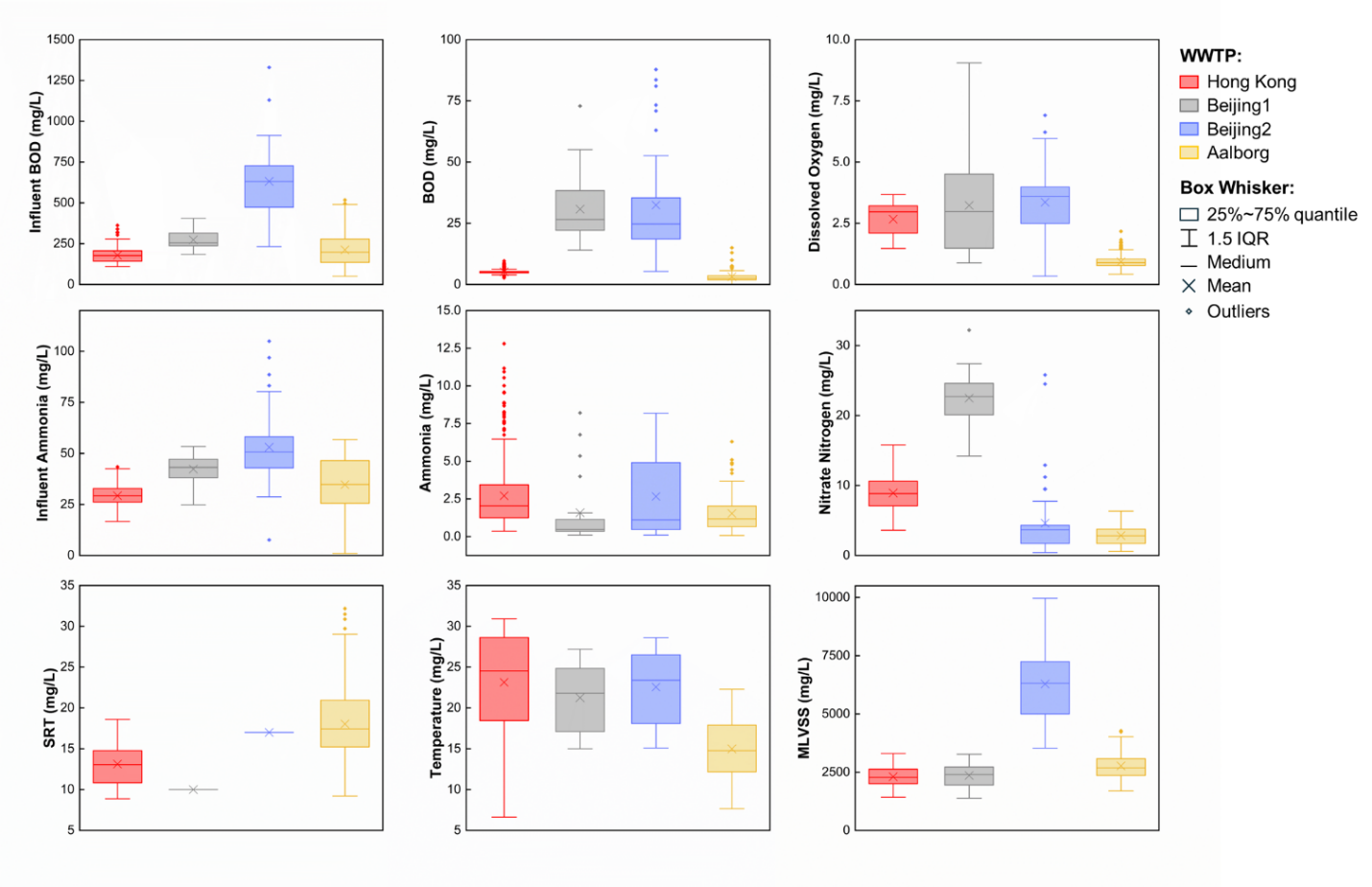

Figure S1. Summary of the environmental factors of the four activated sludge systems. BOD: biochemical oxygen demand. SRT: sludge retention time. MLVSS: mixed liquid volatile suspended solid.

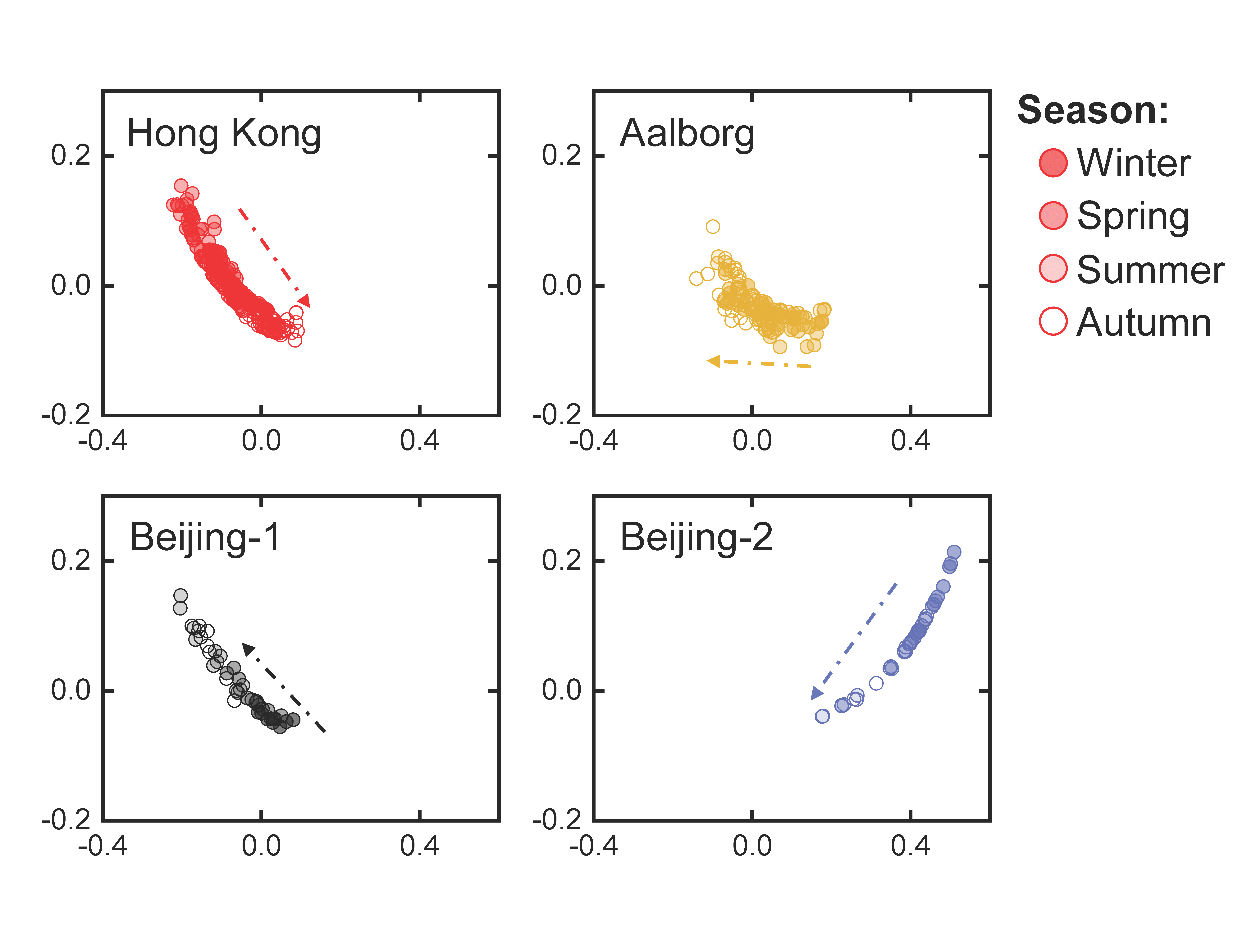

Figure S2. PCoA of the environmental factors of the four activated sludge systems.

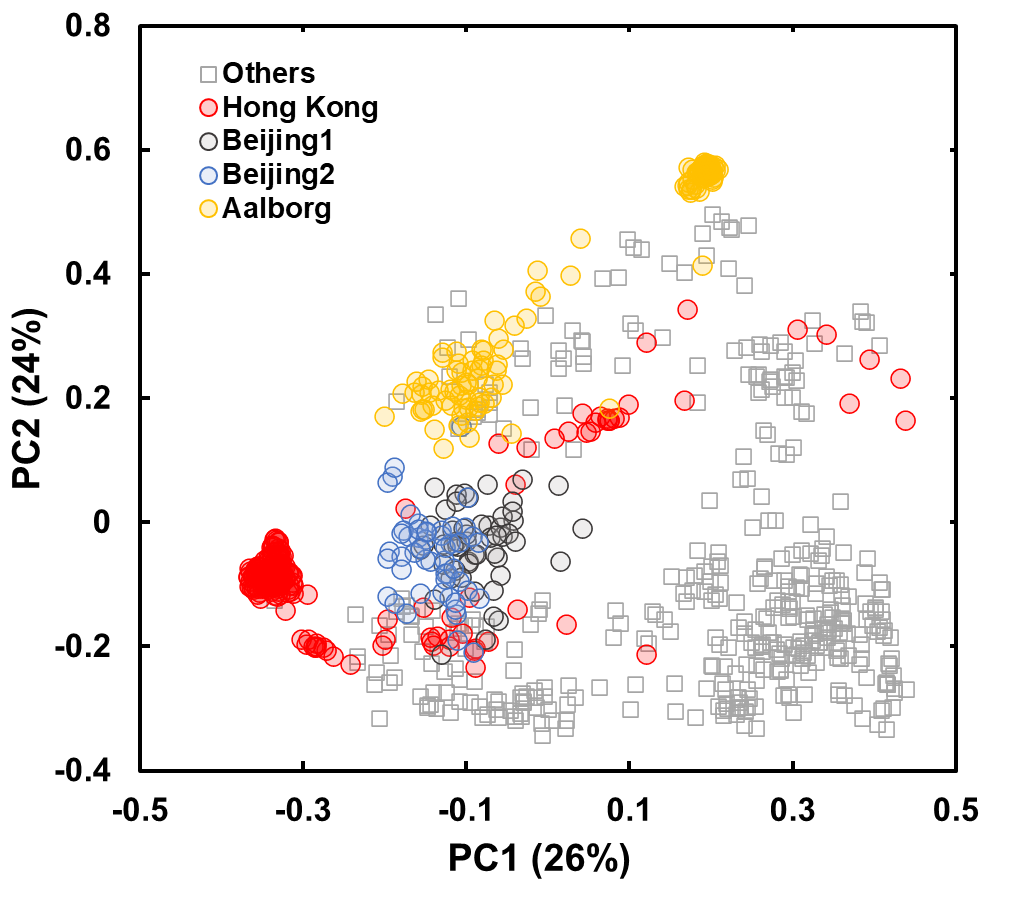

Figure S3. Bray-Curtis distance-based PCoA of activated sludge communities.

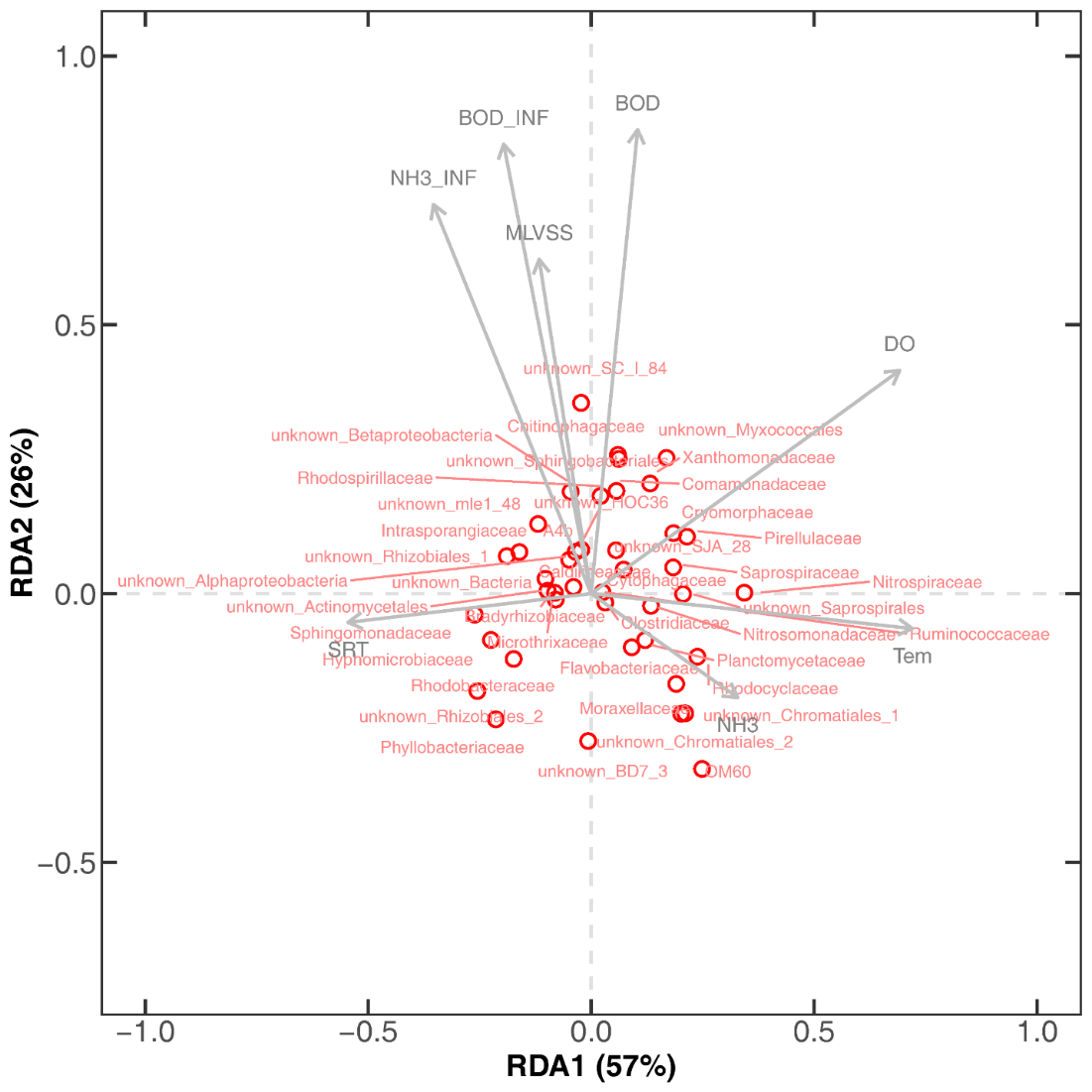

Figure S4. RDA of relative abundance of core populations and environmental factors.

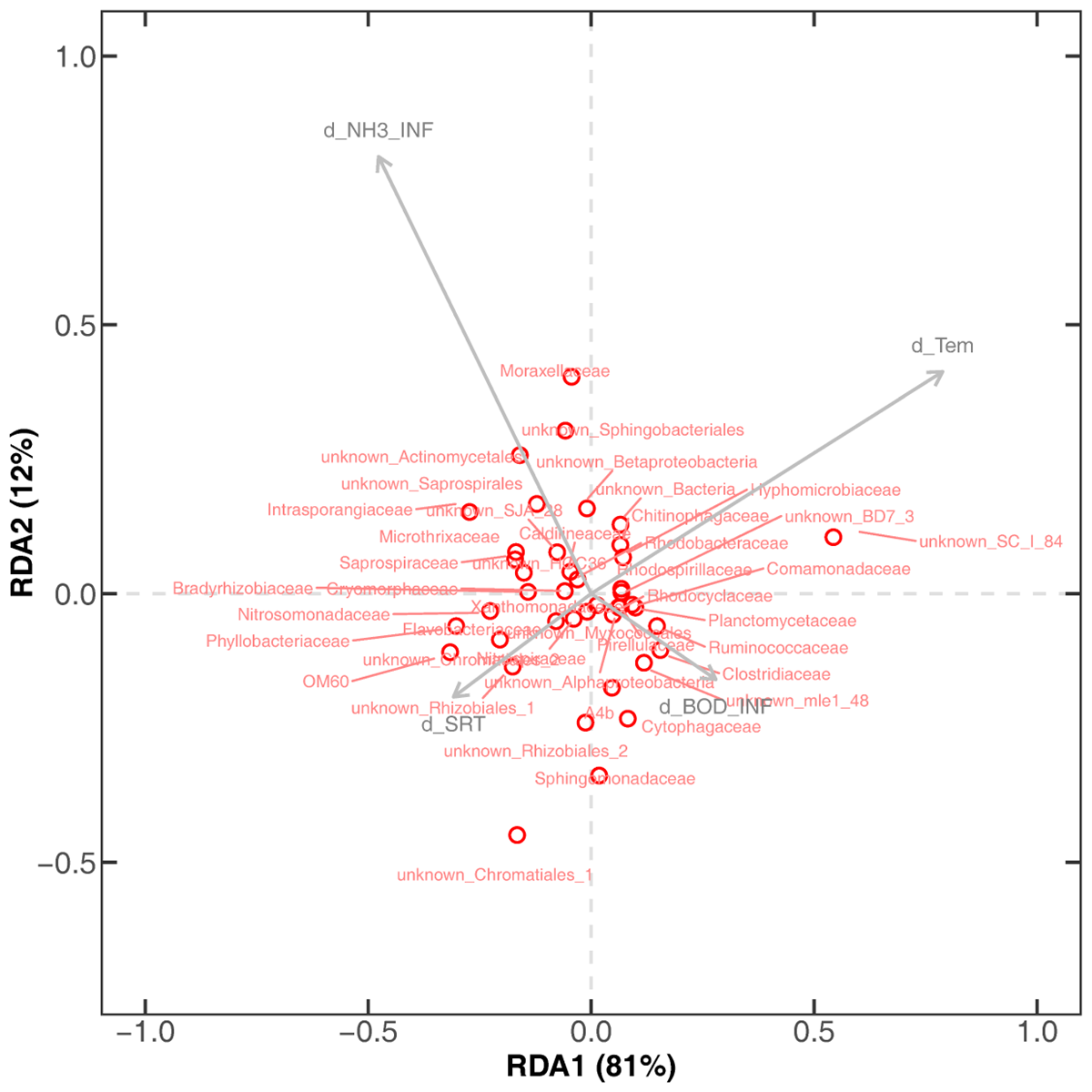

Figure S5. RDA of population dynamics and environmental perturbations.

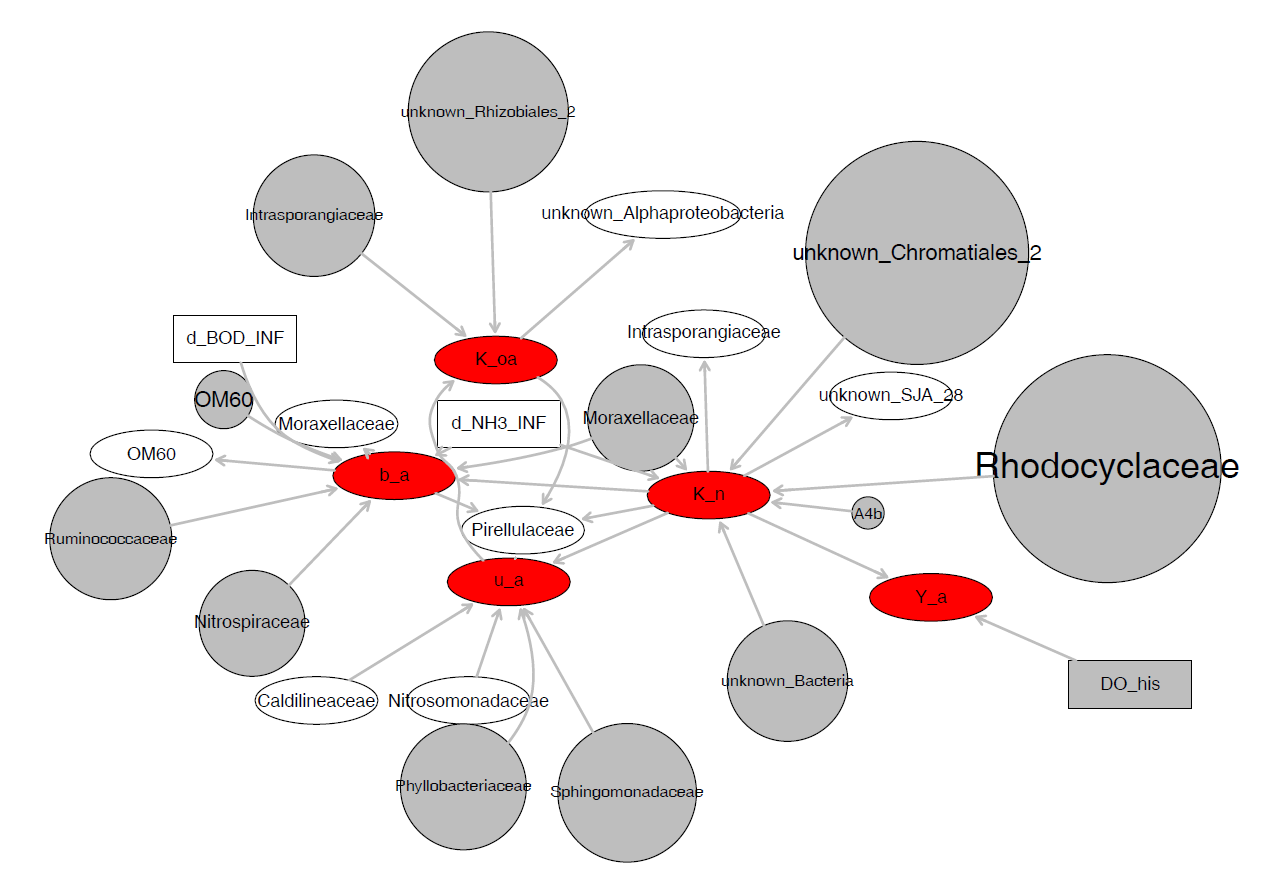
Figure S6. The Bayesian network: related features with the center of autotroph kinetic parameters.

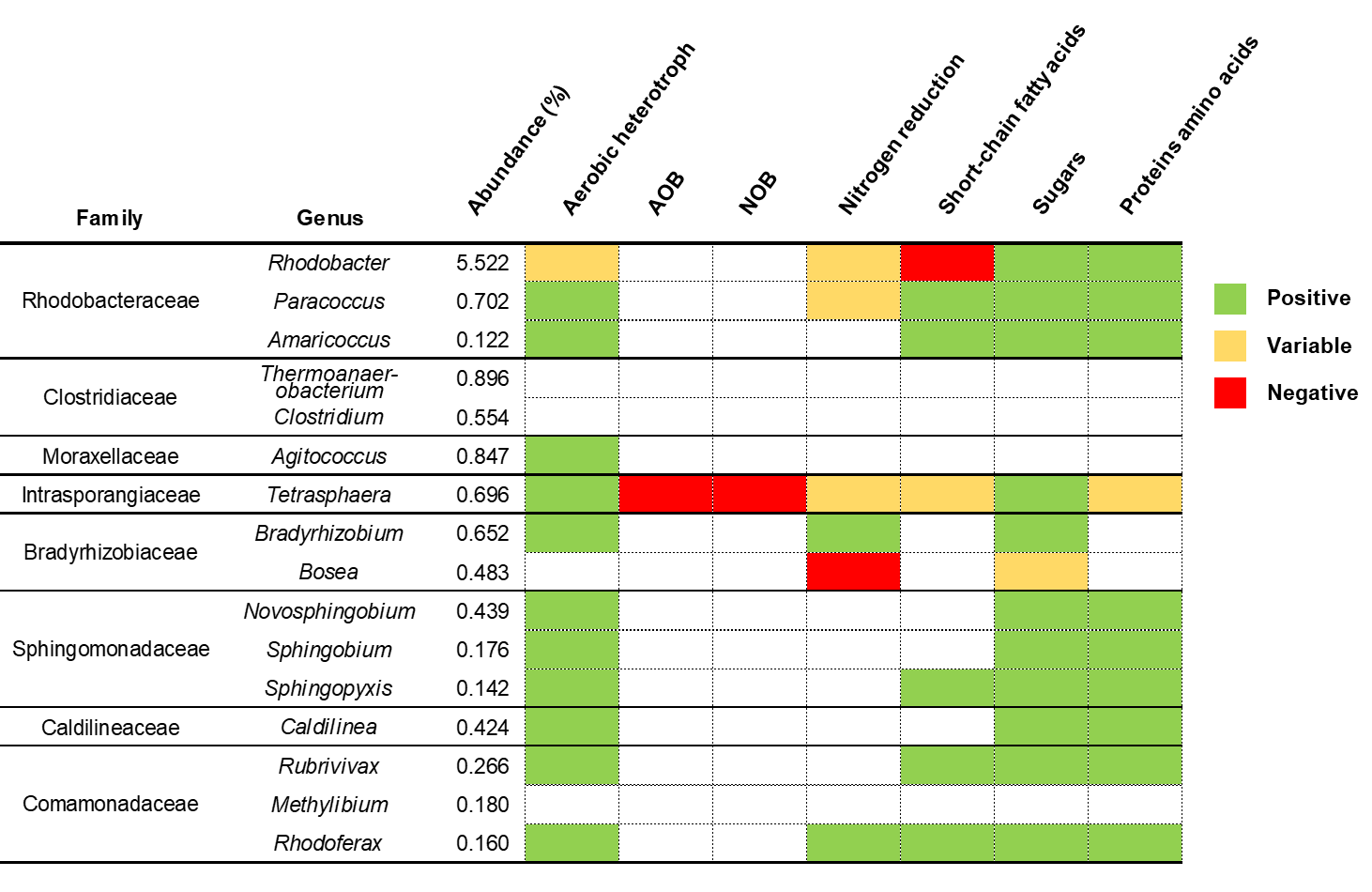

Figure S7. Mapping of the genera from relevant core families in the heterotroph subnetwork to the MiDAS database.

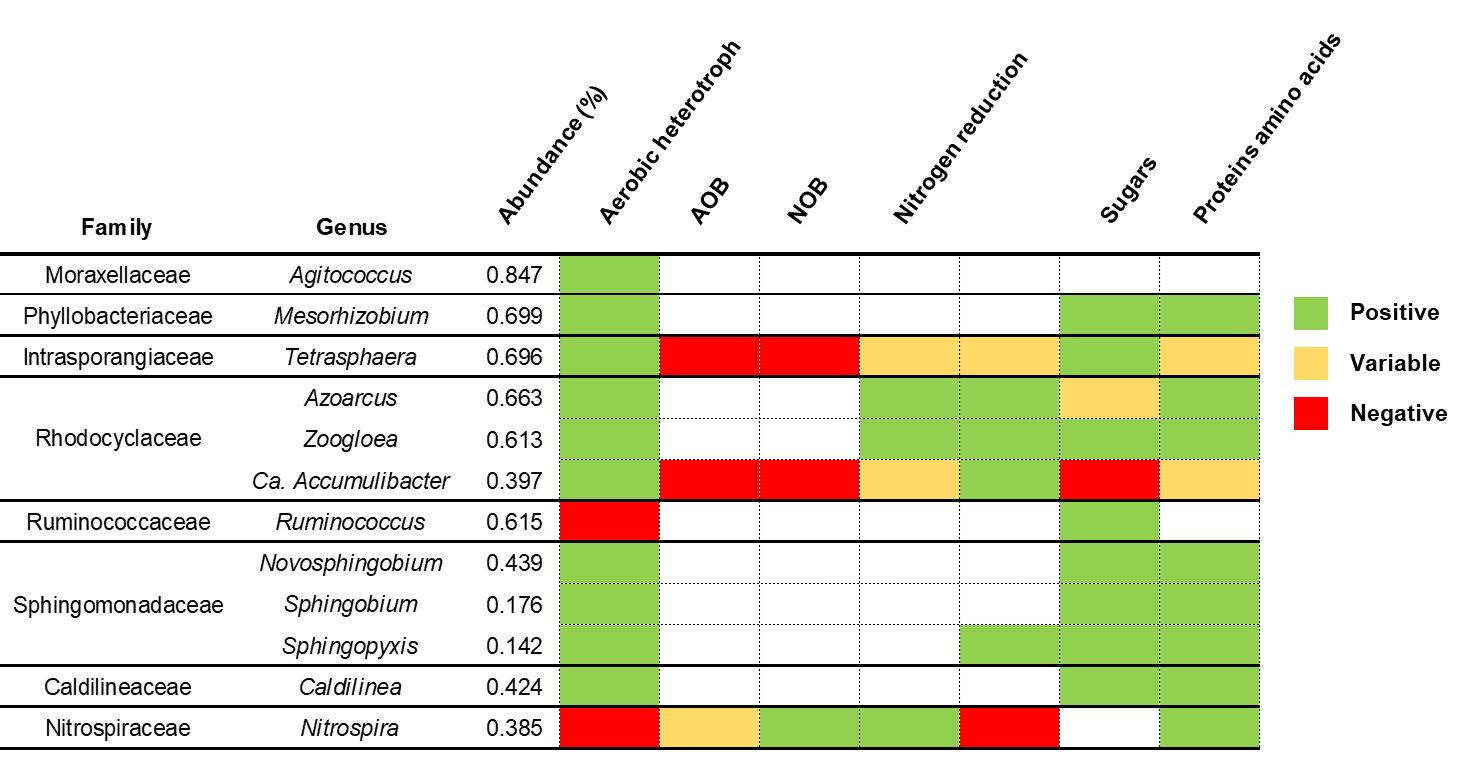

Figure S8. Mapping of the genera from relevant core families in the autotroph subnetwork to the MiDAS database.

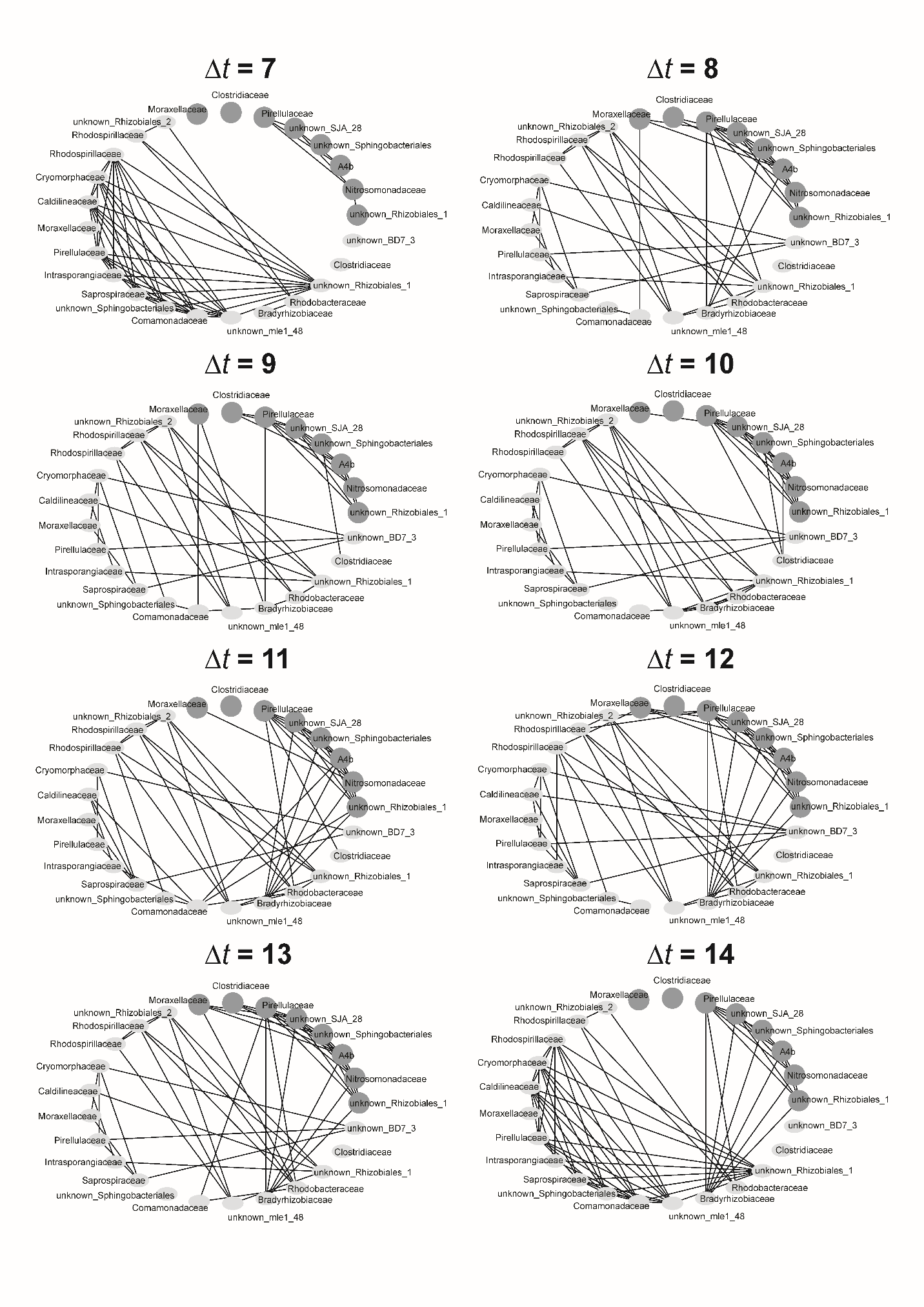
Figure S9. Topological analysis of historical microbial abundance and population dynamics of heterotrophs throughout eight timespans.

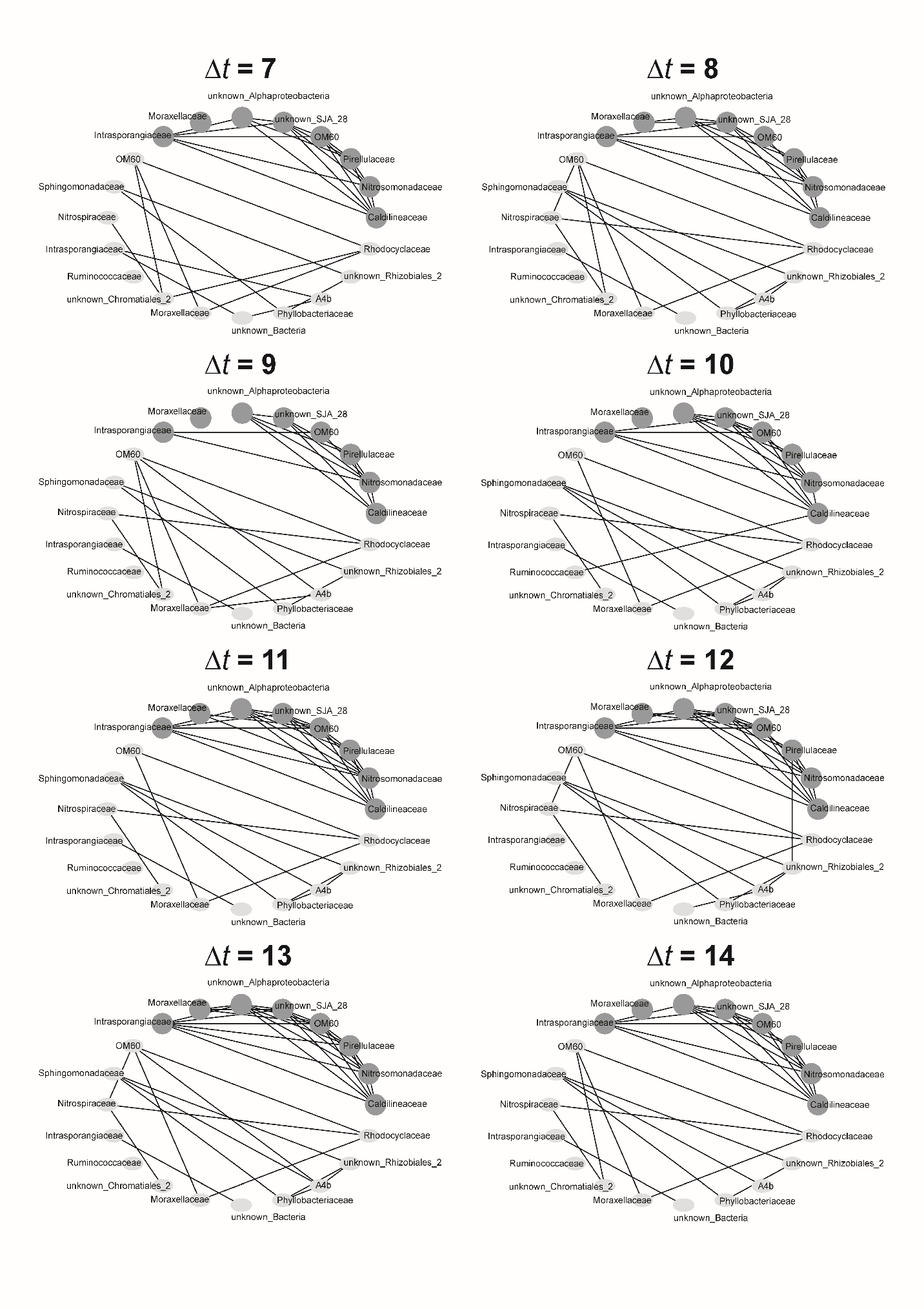
Figure S10. Topological analysis of historical microbial abundance and population dynamics of autotrophs throughout eight timespans.

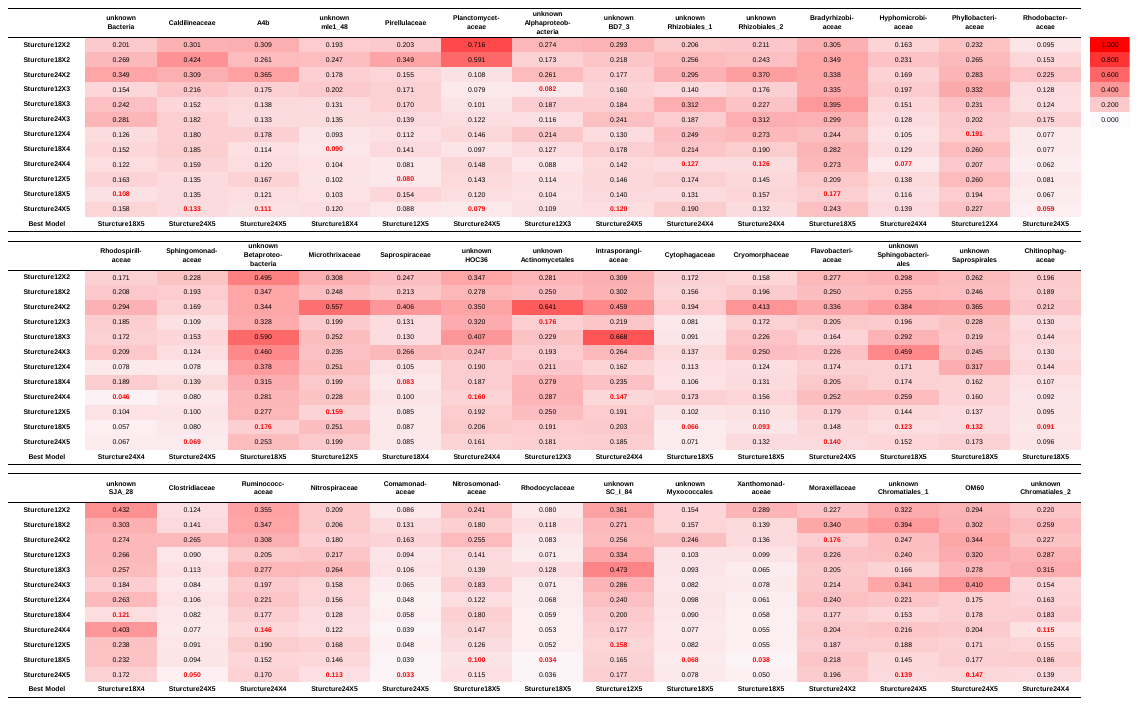
 Figure S11. Averaged RMSE obtained during ANN cross-validation with microbial kinetic parameters as inputs. Numbers in red represent the RMSE of optimal ANNs.

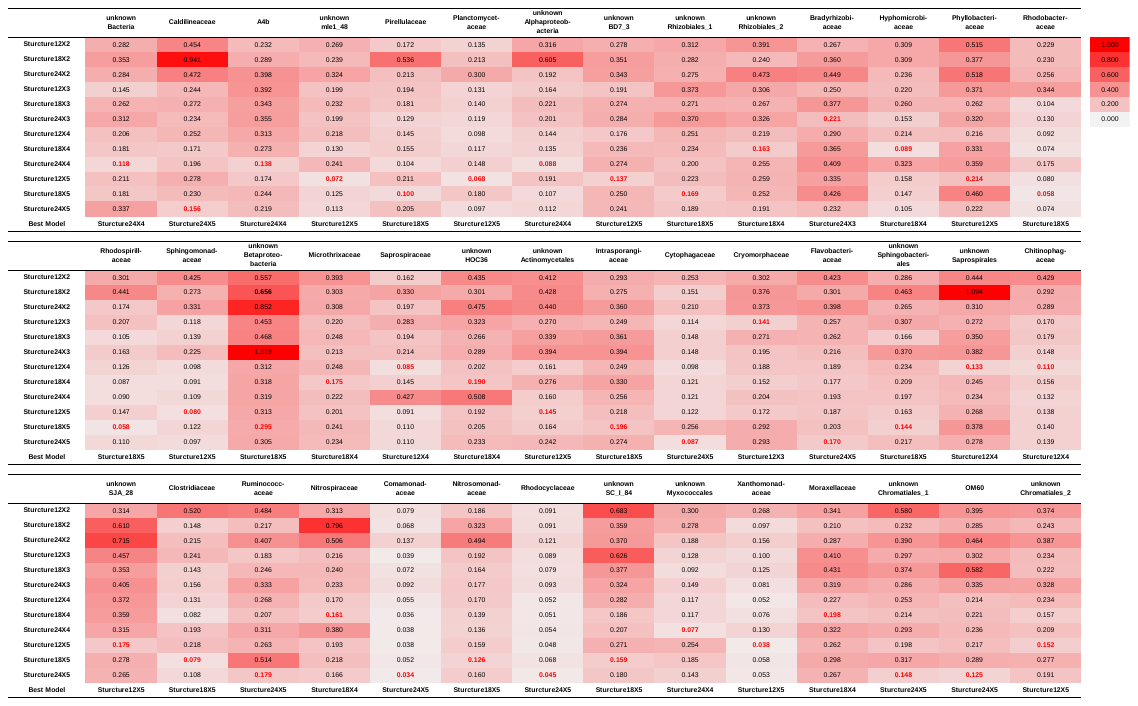

Figure S12. Averaged RMSE obtained during ANN cross-validation with microbial kinetic parameters as inputs. Numbers in red represent the RMSE of optimal ANNs.

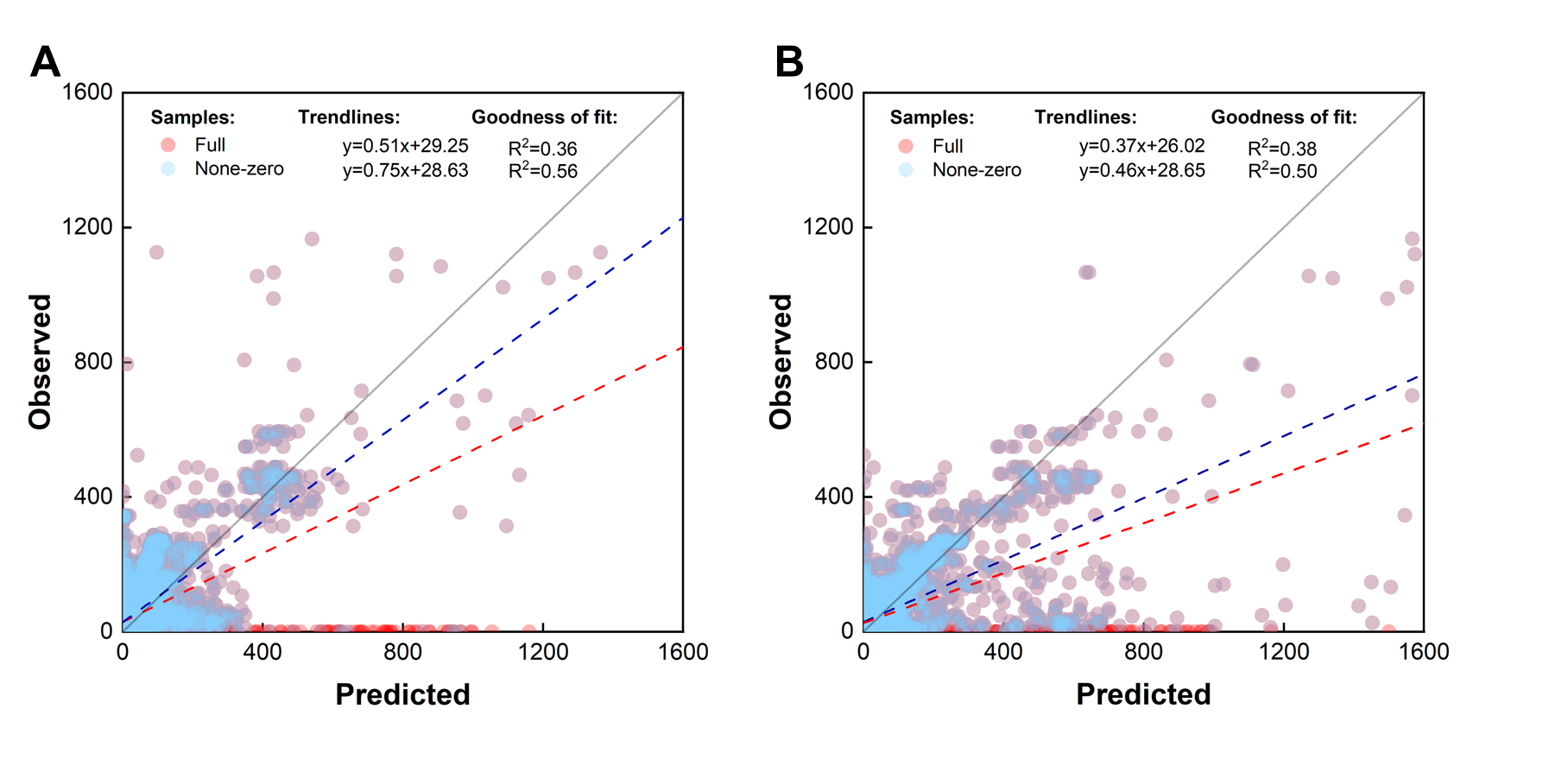

Table S13. Comparison between predicted and observed dynamics of the core populations. (A) Predictions from ANNs with microbial kinetic parameters as inputs. (B) Predictions from ANNs without microbial kinetics parameters.

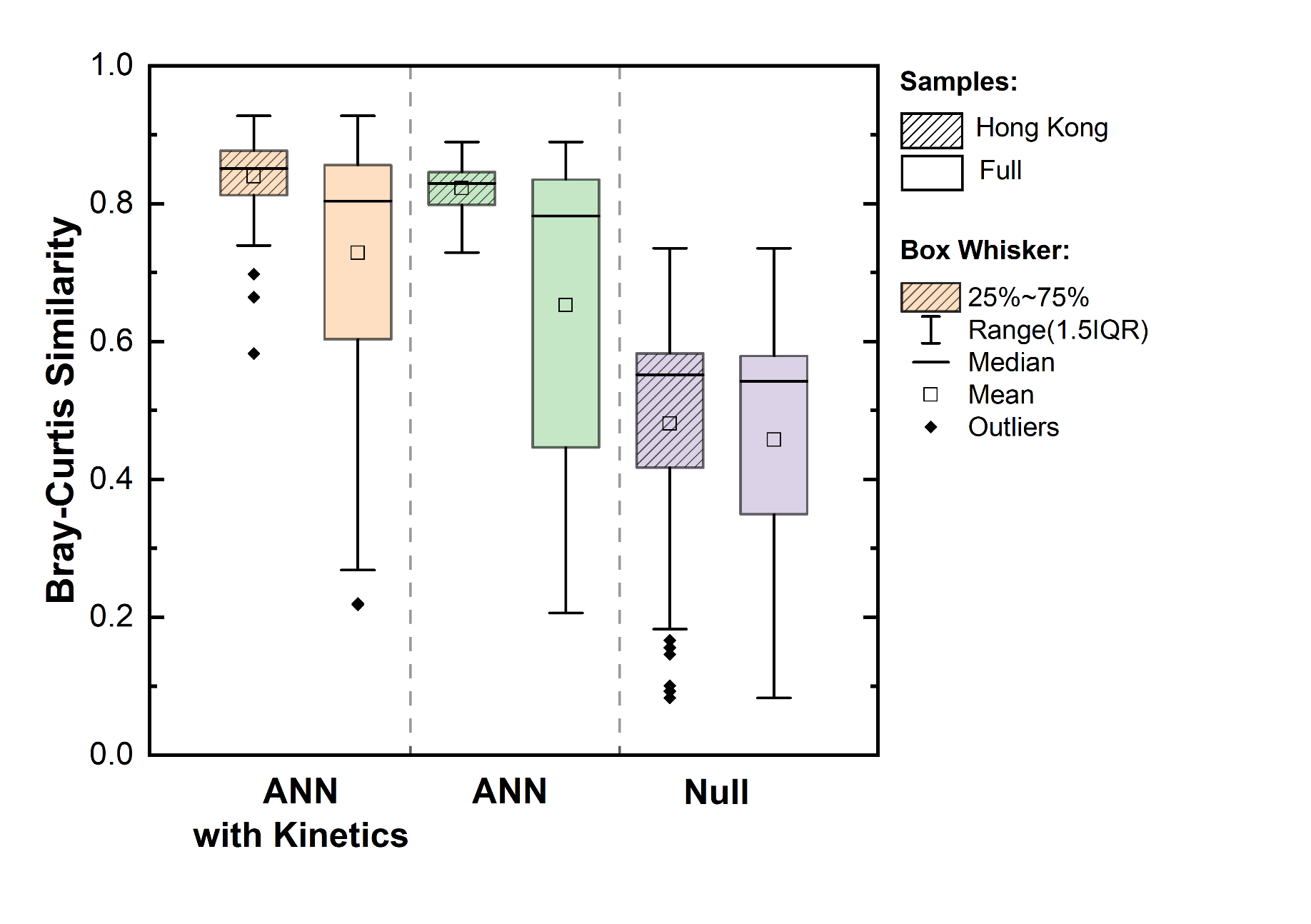

Figure S14. Bray-Curtis Similarity between observed and predicted community structure.
